# Environmental context sculpts spatial and temporal visual processing in thalamus

**DOI:** 10.1101/2024.07.26.605345

**Authors:** Kayla Peelman, Bilal Haider

**Affiliations:** Dept of Biomedical Engineering, Georgia Institute of Technology & Emory University, Atlanta, GA, USA

## Abstract

Behavioral state modulates neural activity throughout the visual system^1–3^. This is largely due to changes in arousal that alter internal brain state^4–10^. Much is known about how these internal factors influence visual processing^7–11^, but comparatively less is known about the role of external environmental contexts^12^. Environmental contexts can promote or prevent certain actions^13^, and it remains unclear if and how this affects visual processing. Here, we addressed this question in the thalamus of awake head-fixed mice while they viewed stimuli but remained stationary in two different environmental contexts: either a cylindrical tube, or a circular running wheel that enabled locomotion. We made silicon probe recordings in the dorsal lateral geniculate nucleus (dLGN) while simultaneously measuring multiple metrics of arousal changes, so that we could control for them across contexts. We found surprising differences in spatial and temporal processing in dLGN across contexts. The wheel context (versus tube) showed elevated baseline activity, and faster but less spatially selective visual responses; however, these visual processing differences disappeared if the wheel no longer enabled locomotion. Our results reveal an unexpected influence of the physical environmental context on fundamental aspects of early visual processing, even in otherwise identical states of alertness and stillness.

## Results and Discussion

Our goal was to determine whether the quality of visual processing in the dLGN depends upon external environmental context, independent from effects driven by differences in arousal. (Fig. 1A, B). Our main measure of arousal level was cortical local field potential (LFP) power in low frequencies (1 – 4 Hz). Although this is just one of many potential metrics for assessing arousal level, it proved highly effective in capturing arousal fluctuations that were correlated with many other internal and external variables, as will be detailed below. Therefore, in this study, increasing arousal is defined by reductions in low frequency cortical LFP power (Fig. 1E).

**Figure 1.**
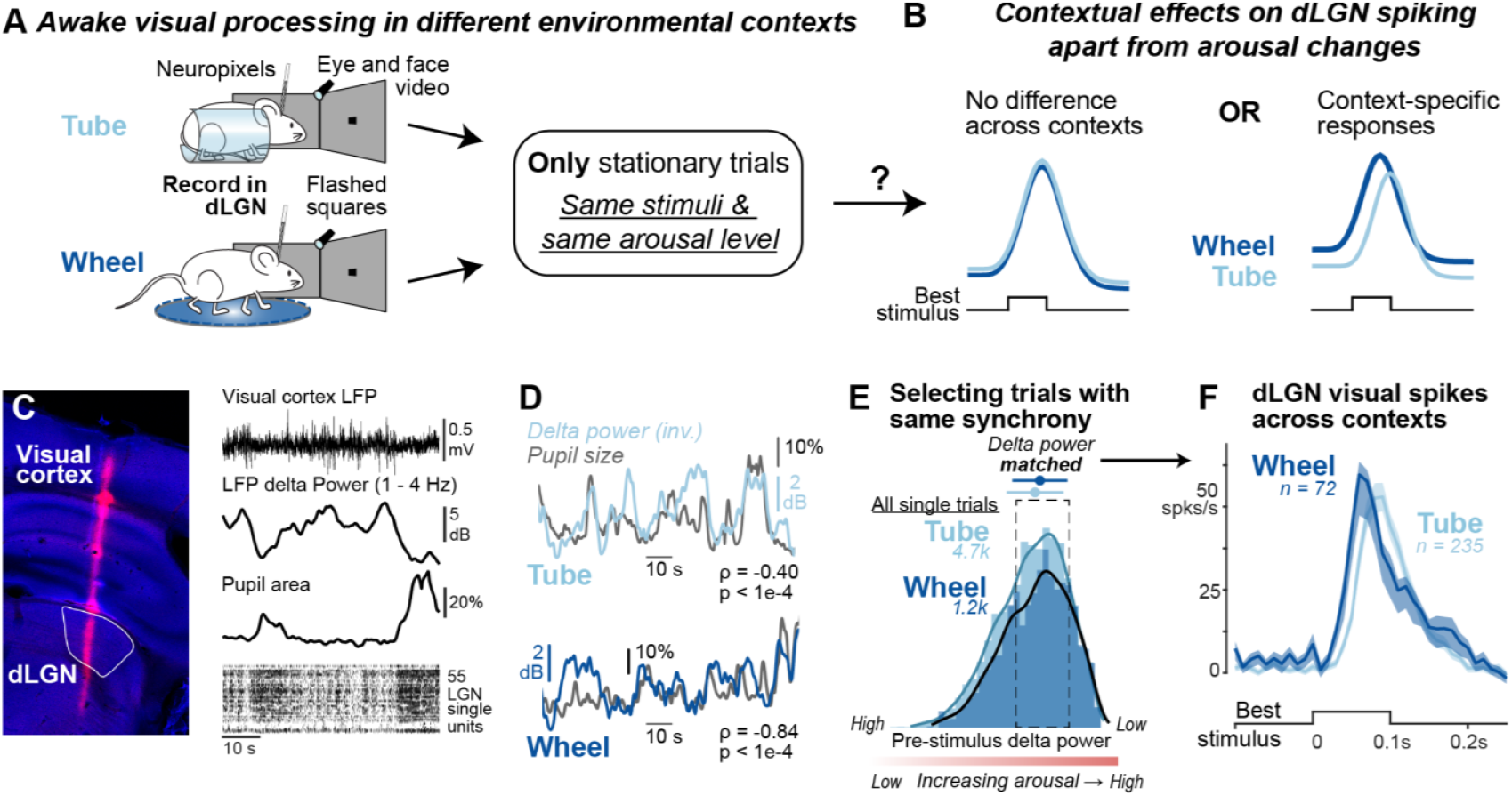
Measuring awake dLGN responses and arousal level in two different environmental contexts. **A**. Awake, head-fixed mice were placed either in a tube (N=9 mice) or on circular treadmill (N=5 mice) while viewing full-contrast black or white squares (7°) presented one at a time for 0.1s in random locations across the visual hemifield. Neuropixels 1.0 probes simultaneously sampled neural activity across visual cortex and the dorsal lateral geniculate nucleus (dLGN). We examined only stationary trials on the wheel (72% of total, see Fig. S1) to isolate effects of context. **B**. Schematic of two hypotheses for contextual effects. *Left*, visual responses in dLGN show no differences across context when arousal is the same. *Right*, dLGN responses show context-specificity even when arousal is the same. **C**.*Left*, histological confirmation of probe track passing through visual cortex and targeting dLGN. *Right*, local field potential (LFP) and delta (1-4 Hz) power was measured in visual cortex while single units were isolated in dLGN. Simultaneous high-speed video captured pupil and periocular motion. **D**. Example traces of delta power (inverted) and pupil size across contexts. (Pearson correlation in tube ρ = −0.40, *p* < 0.001; wheel ρ = −0.84, *p* < 0.001). **E**. Distribution of delta power preceding all analyzed visual stimulus trials in tube (light blue; 4734 trials in central bin) and wheel (dark blue; 1178 trials). Note that delta power axis is reversed (arranged high to low), so that arousal level increases from left to right. The matched delta power range (dashed box) displays the median ± 1 MAD of the normalized pre-stimulus delta power (reflecting similar cortical synchrony) when combined for both tube and wheel stationary trials (wheel distribution: 0.060 ± 0.026 normalized delta power values; tube distribution 0.062 ± 0.027; p = 5.1196e-05, Kolmogorov–Smirnov test). **F**. dLGN neuron spike responses to the preferred stimulus in the receptive field across contexts (tube, n = 235 neurons; wheel n = 72 neurons).

We tested the hypothesis that context itself engenders different qualities to visual processing, even during identical states of arousal; an alternative hypothesis is that visual processing is largely explained by arousal state, regardless of context. To test these hypotheses, we placed awake, head-fixed mice in two distinct environmental contexts: a semi-enclosed plastic cylinder (“Tube” group; N = 9 mice), or on a circular treadmill in which mice were free to locomote or remain stationary (“Wheel” group; N = 5 mice). Experiments were often performed in both contexts in the same mice (5/9 mice were recorded from in both contexts). When performing experiments in these 5 mice, we randomized which context the animal would be in that day, so there was no systematic bias in the order of exposure to wheel versus tube. On the wheel, we verified well-known effects of locomotion-induced arousal changes on dLGN visual processing (Fig. S1), indicating the wheel context captures a behavioral repertoire investigated in many prior studies^9,14^. Here, we focus on the most frequent behavioral epochs (when mice were stationary on the wheel; 72% of time; Fig. S1J), and compared these with mice stationary in the tube. This permits comparison of visual processing in two distinct contexts, and on trials with similar arousal levels (explained below), free from influences of ongoing locomotion itself (Fig. 1A-B).

We simultaneously recorded spiking in dLGN and local field potential (LFP) activity from higher visual areas of cortex using Neuropixels 1.0, along with high-speed video of the face and eye (Fig. 1C). To assess contextual effects on spatial and temporal receptive field (RF) responses of single neurons, we presented small, brief black and white squares (7°; 0.1s duration) in randomized locations throughout the contralateral visual hemifield. We identified trials with the preferred stimuli for each dLGN neuron (stimulus color and location eliciting maximal response; Methods), and responses to these stimuli are the main topic of this report.

Cortical LFP delta power levels during stationary trials were highly overlapping in the two contexts. We categorized arousal levels using the visual cortical LFP power (1-4 Hz, or “delta”), a “gold standard” electrophysiological signature of arousal that is tightly synchronized with thalamic activity^15,16^, and changes in pupil size^6^, thus providing a high temporal resolution measure of trial-by-trial arousal levels across the visual thalamocortical pathway. Cortical delta power is also widely used to categorize “synchronized” and “desynchronized” states in cortex that are tightly linked to low and high arousal, respectively^17–19^. In both contexts, visual cortical delta power was significantly anti-correlated with pupil size, another well-known indicator of arousal (Fig. 1D; delta power inverted for display). Importantly, although pupil size varied with arousal level, the effects we detail in the following sections could not be explained simply by more light entering the pupil (Fig. S2; controls in full darkness and with pharmacologically dilated pupil). The trial-by-trial distribution of cortical delta power (reflective of cortical synchrony) was highly overlapping across both contexts (Fig. 1E), but significantly different across the entire distributions. We therefore focused on the central 55% of trials surrounding the modal delta power level to examine the impact of context itself – apart from arousal level – on dLGN visual responses.

To our surprise, in these carefully matched conditions, the overall dLGN spiking response to stimulation of the RF showed context specificity, with visibly faster and slightly stronger visual responses in stationary epochs on the wheel versus tube (Fig. 1F; wheel peak response is ~10 ms faster and evokes 4.8 more spikes per second relative to the tube). In the following sections, we quantify in detail how context governs the amplitude, timing, and selectivity of these responses.

### Context adjusts visual response timing and baseline activity in dLGN

We first found that context influenced both baseline activity and visual response timing (Fig. 2A).

**Figure 2.**
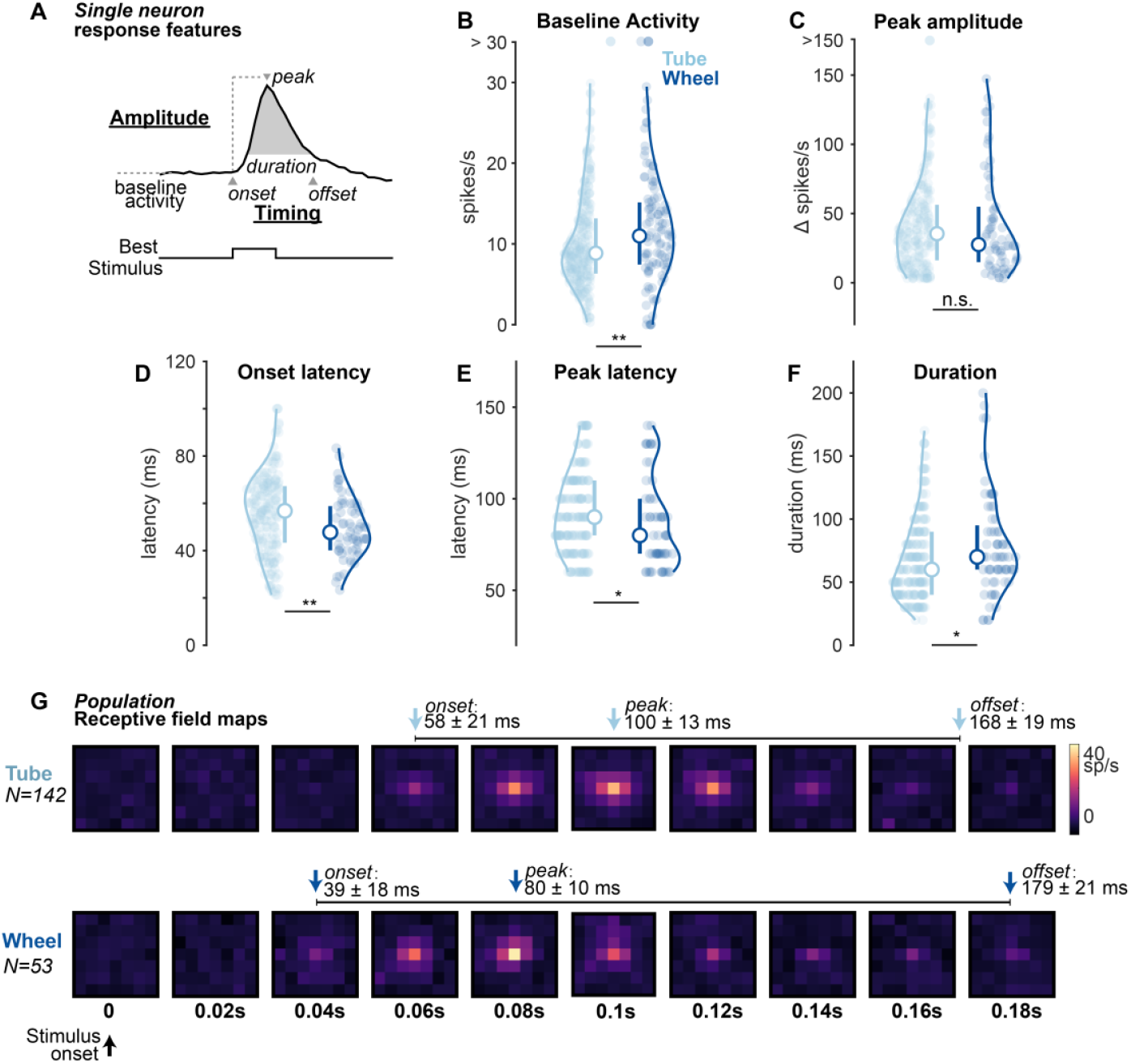
Environmental context influences visual response timing in dLGN. **A**. Schematic for quantification of visual response features (amplitude and timing) in dLGN across contexts. **B**. Baseline activity was significantly elevated in the wheel context (wheel: 11.4 ± 4.1 spikes/s, n = 107; tube: 8.9 ± 3.4 spikes/s, n=276; *p* = 0.0018; Wilcoxon one-sided rank-sum test, Median ± IQR/2; same statistics throughout figure). Baseline calculated in 0.1s window prior to stimulus onset. **C**. Baseline-subtracted peak visual response similar across contexts (wheel: Δ23.3 ± 18.9 spikes/s, n=72; tube: Δ30.7 ± 21.3 spikes/s, n=235; *p* = 0.088). **D**. Onset latency significantly faster on the wheel (wheel: 47 ± 19 ms, tube: 57 ± 24 ms; *p* = 0.003). **E**. Time to peak response significantly faster on the wheel (wheel: 87 ± 15 ms, tube: 93 ± 15 ms; *p* = 0.020) **F**. Response duration significantly longer on wheel (wheel: 70 ± 25 ms, tube: 60 ± 20 ms; *p* = 0.025). **G**. Population receptive field maps across time aligned to the center of each neuron’s receptive field (top, tube n = 142 units; bottom, wheel n = 53). Population response onset and peak significantly faster on wheel (onsets: wheel, 39 ± 18; tube, 58 ± 21 ms; *p* = 0.002; peaks: wheel, 80 ± 10 ms; tube, 100 ± 13 ms; *p* = 0.044; Mean ± SD, permutation test; Methods). Response offset and overall duration (where activity was above 25% of the maximum response amplitude) not significantly different (tube 168 ± 19 ms, wheel 179 ± 2 ms, *p* = 0.360).

Baseline activity was significantly greater on the wheel relative to the tube (Fig. 2B; wheel: 11.4 ± 4.1 spikes/s, tube: 8.9 ± 3.4, *p* = 0.0018). Accordingly, both spontaneous and visually evoked bursting (Methods) was significantly reduced in the wheel compared to the tube (spontaneous p = 0.007; visually evoked p = 9.93e-07, Fig. S2A, B). After accounting for these baseline activity differences, the peak visual response was not significantly different across contexts (Fig. 2C; wheel: Δ23.3 ± 18.9 spikes/s, tube: Δ 30.7 ± 21.3, *p* = 0.088).

However, visual response onset and peak response latencies were both significantly faster by ~10 ms on the wheel (Fig. 2D,E; onset latency: wheel: 47 ± 19 ms, tube: 57 ± 24 ms; *p* = 0.003; time to peak: wheel: 87 ± 15 ms, tube: 93 ± 15 ms; *p* = 0.020). The response duration was also significantly prolonged on the wheel (Fig. 2F; wheel: 70 ± 25 ms, tube: 60 ± 20 ms; *p* = 0.025). Overall, across all stationary epochs with highly overlapping arousal levels, visual responses were faster and more prolonged on the wheel than the tube.

To ensure that these timing differences were not due to individual sessions, neurons, or mice, we generated population receptive field maps from many combinations of neural ensembles recorded in each context (Fig. 2G; Methods). These surrogate populations were subsampled randomly from the entire recorded database to match the number of recordings in each context (17 in tube, 6 on wheel). Similar to the single neuron responses, we again found that the population response onset and peak latencies were faster on the wheel, (onsets: wheel, 39 ± 18; tube, 58 ± 21 ms; *p* = 0.002; peaks: wheel, 80 ± 10 ms; tube, 100 ± 13 ms; *p* = 0.044), but duration was comparable. This resampling approach ensures that the data analysis was not driven by specific recording sessions or neurons, reduces potential biases, and provides a more robust comparison of response properties across contexts. These findings on response timing also persisted when we separately examined ON and OFF cells (ON cells: wheel, *p* = 0.0475; OFF cells: *p* = 0.0184), when we equalized samples across contexts (n=100 resamples of matched neuron populations between tube and wheel, onset latency p < 0.05 for 100/100 resamples), and across individual animals (Fig. S7). The differences were also not explained by RF location biases across contexts (Methods). Lastly, using a multi-dimensional classification of single-trial arousal (with either internal brain signals or external bodily signals) did not appreciably change the results (Fig. S4).

### Increasing arousal elevates baseline activity and improves response timing

The results so far demonstrate effects of context on visual processing when arousal levels were in an overlapping range. Does context also shape the way that changes in arousal level (i.e., increasing from low to high) affect dLGN visual responses? We addressed this by splitting the distribution of trials examined thus far into five equally sized bins, and arranged them from low to high arousal (Fig. 3A; Methods). To ensure that the definition of these bins remained independent of context, we combined all data across all recordings in both contexts and defined the bins according to the overall distribution of pre-stimulus cortical delta power. These bin definitions were then applied to each individual context for all comparisons (Fig. 3B, C). We note that the highest arousal bins in stationary trials were distinct from— but overlapping with— the highest arousal bins observed during locomotion (Fig. S1D).

**Figure 3.**
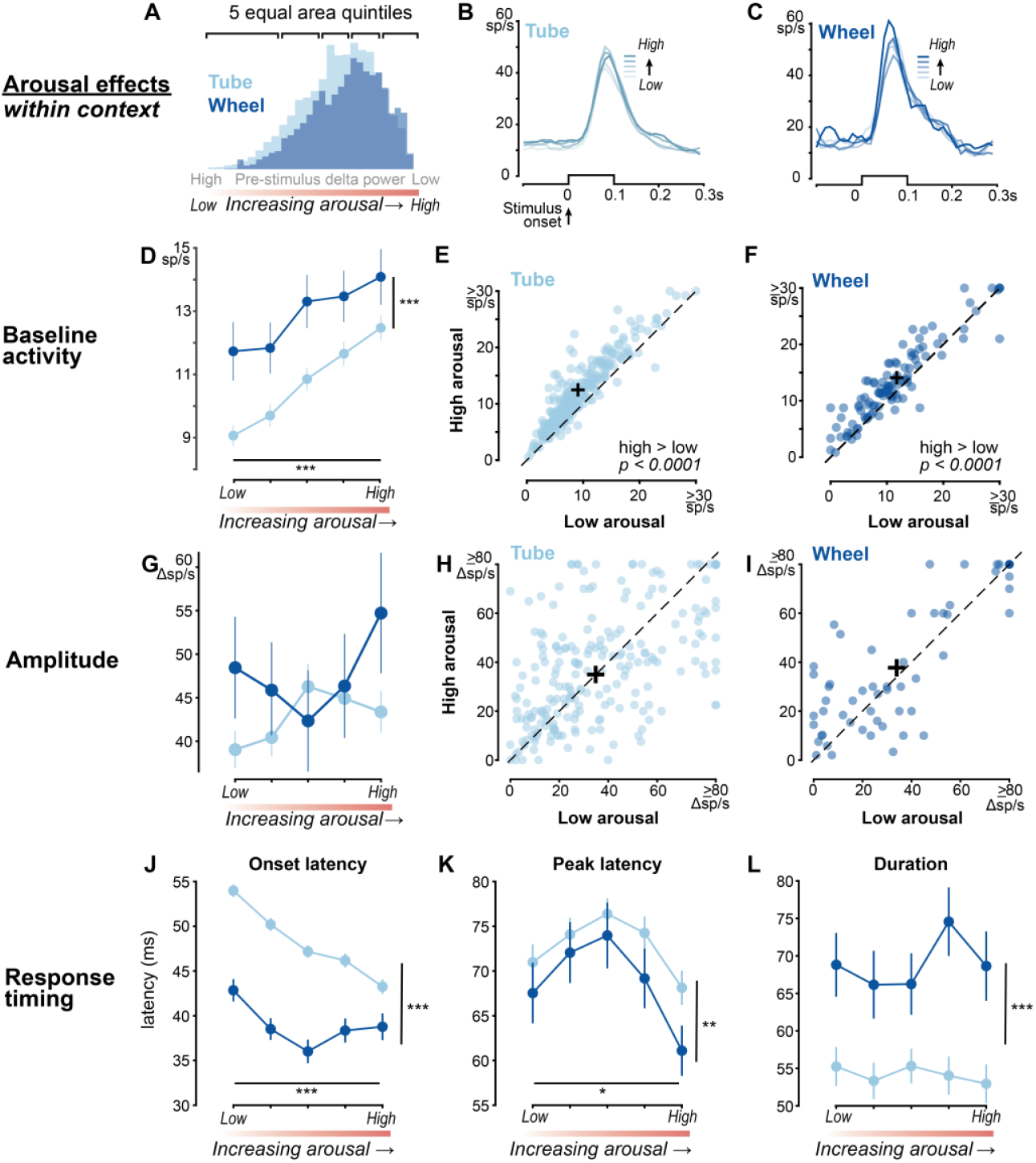
Impact of context on arousal-mediated changes of visual responses in dLGN. **A**. Pre-stimulus delta power distributions for tube (light) and wheel (dark) contexts divided into five equally sized bins to examine responses across arousal level. **B**. Average responses to best stimulus in the RF for tube (n=223 neurons), sorted by increasing arousal levels (line opacity). **C**. Same as in B, but on the wheel (n=63 neurons). **D**. Significant effect of context (*p* = 2.2e-9) and arousal (*p* = 1.9e-6) on baseline activity (two-way ANOVA throughout figure unless indicated). Baseline activity increases with increasing arousal in both contexts (Spearman correlation, tube r = 1.00, *p* = 0.0167; wheel r = 1.0, *p* = 0.0167). **E**. Baseline activity significantly greater at high versus low arousal in tube (low: 9.1 ± 0.3 spikes/s; high: 12.5 ± 0.4; *p* = 4.5e-41, Mean ± SEM; sign rank test). **F**. Same as E, for wheel (low: 11.7 ± 0.9 spikes/s, high: 14.1 ± 0.9; *p* = 3.7e-8, sign rank test). **G**. Baseline-subtracted visual response amplitude not significantly different with arousal level (*p* = 0.59) or context (*p* = 0.057). **H**. Baseline-subtracted visual response amplitude similar for low versus high arousal levels in the tube (low: Δ39.0 ± 2.4 spikes/s, high: Δ 43.5 ± 2.6; *p* = 0.260, sign rank test). **I**.Same as H, for wheel (low: Δ48.4 ± 6.0 spikes/s, high: Δ54.7 ± 7.0 spikes/s; *p* = 0.083, sign rank test). **J**. Faster onset latency with increasing arousal (*p* = 2.92e-5) and is overall fastest on the wheel (*p* = 3.2e-19). **K**. Time to peak response is overall faster in the wheel context (*p* = 0.02). **L**. Response duration) is longer in the wheel context (*p* = 4.0e-9).

We found that the influence of arousal level on dLGN activity exhibited similar directionality (i.e., increasing arousal drove increasing firing), but with significantly different set points across contexts. First, as expected, we found that increasing arousal level significantly elevated baseline activity for both contexts (Fig. 3D; Spearman correlation, tube r = 1.00, *p* = 0.0167; wheel r = 1.0, *p* = 0.0167). This relationship was highly correlated on both the wheel and the tube, such that baseline activity was significantly greater at highest versus lowest arousal bins (Fig. 3E, F; 12.5 ± 0.4 spikes/s versus 9.1 ± 0.3; *p* = 4.5e-41). However, regardless of arousal level, baseline activity was overall significantly greater on the wheel versus tube (Fig. 3D; p = 2.2e-9). Second, we found that increasing arousal level did not have a significant effect on visual response amplitude (Fig. 3G; p = 0.590), with no clear difference on visual responses between highest and lowest arousal groups within context (Fig. 3H, I; tube: p = 0.26, wheel p = 0.083). Lastly, we found that increasing arousal significantly accelerated visual response onset latencies within context, (Fig. 3J; p = 2.92e-5), and these were again overall fastest on the wheel (p = 3.2e-19). Peak response latencies showed a similar form of arousal-induced modulation across contexts, and again overall these were fastest on the wheel (Fig. 3K; p = 0.020). Response duration was significantly prolonged on the wheel (p = 4.0e-9), but invariant to increasing arousal levels within context (Fig. 3L; p = 0.8124). These findings on the relationship between increasing arousal and context were highly similar when we instead used other well-known measures of arousal to classify the trials (such as pupil size, narrowband gamma (NBG) power (50-70 Hz), facial motion energy; Fig. S3), or combinations of these multiple variables (Fig. S4). Taken together, our results show significant arousal-induced changes to baseline activity and visual response timing in dLGN even in stationary epochs, but with context-specific expression of these relationships.

### Both context and arousal impact receptive field size & selectivity

Thus far, we have examined contextual effects on visual response amplitude and timing. Does context affect visual spatial resolution? First, we looked at how receptive field (RF) size and RF selectivity (Fig. 4A) differed across contexts at matched levels of arousal (Fig. 4B, C, example units; same analysis as in Figs. 1 and 2). We found that, overall, RF size was significantly smaller on the wheel (Fig. 4D; wheel 12 ± 7°, tube: 19 ± 7°, *p* = 7.9e-3). Overall RF selectivity (center versus background firing) was not different across the two contexts (Fig. 4E; tube: 0.6 ± 0.2; wheel: 0.5 ± 0.2, *p* = 0.260). We then examined how increasing arousal influenced these RF properties. In both contexts, as arousal increased so did RF size (Fig. 4F; *p* = 0.0001), and this significantly reduced RF selectivity (Fig. 4G; *p* = 0.0006). Furthermore, when examining how changes in arousal level effect RF size and selectivity, there emerged a significant effect of context: RFs were smaller (*p* = 0.007) but significantly less selective (relative to background activity) on the wheel (*p* = 0.033). Thus, in both contexts, increasing arousal uniformly broadened RFs and lowered spatial selectivity. However, there remained a more specific effect of context on arousal-induced changes to RF properties: they were significantly smaller but less selective on the wheel versus the tube.

**Figure 4.**
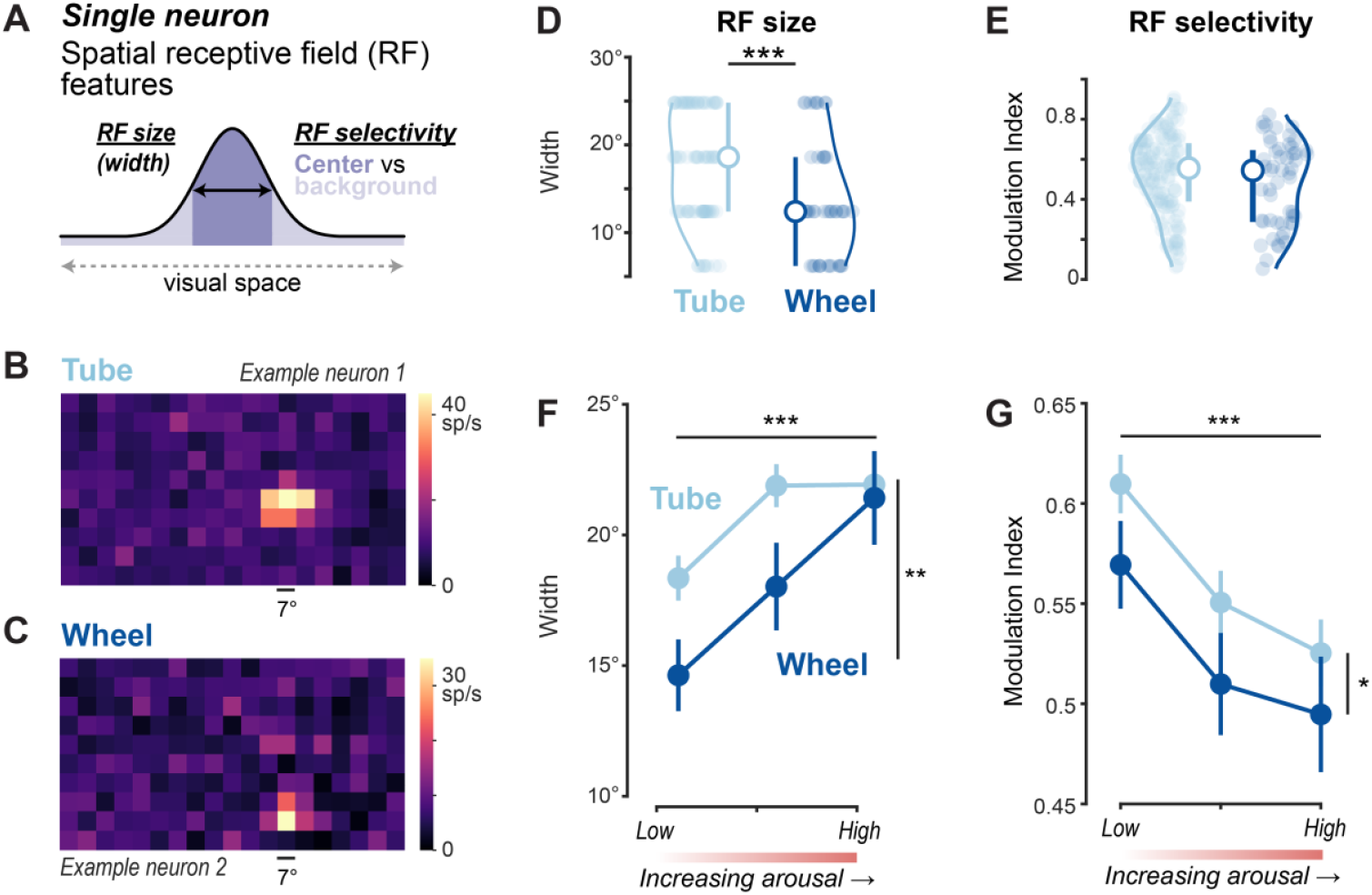
Context-specific effects on dLGN spatial RFs. **A**. Schematic showing estimation of spatial receptive field (RF) size. Width quantified as the extent of contiguous pixels on RF map that evoke ≥ 50% of the maximum amplitude. RF selectivity is quantified with a modulation index (MI) between the center and background [MI = (center − background) / (center + background)]. Center defined as the width at 50% maximum firing rate. No eye movements in any trials analyzed in this figure (Methods). **B**.Example neuron RF map in the tube. **C**.Example neuron RF map on the wheel. **D**.At matched levels of arousal (as in Fig. 1E-F, Fig. 2), RF size is significantly smaller in the wheel (wheel 12 ± 7°, n=43; tube: 19 ± 7°, n = 90; *p* = 7.9e-3; Median ± IQR/2, Wilcoxon one-sided rank-sum test). **E**.At matched levels of arousal, RF selectivity is similar between the tube and wheel contexts (tube: 0.6 ± 0.2; wheel: 0.5 ± 0.2, *p* = 0.260; Median ± IQR/2, Wilcoxon one-sided rank-sum test). **F**. RF sizes significantly smaller on the wheel (*p* = 0.007). On both the tube and wheel, RF size significantly broadens with increasing arousal level (*p =* 0.0001; Mean ± SEM, two-way ANOVA). **G**. RF selectivity is overall higher in the tube (*p* = 0.033) and selectivity worsens with increasing arousal across contexts (*p* = 0.0006; Mean ± SEM, two-way ANOVA).

Finally, are all these contextual differences in visual processing during stationary trials due to the potential for locomotion, or are they perhaps explainable by differences in ongoing postural tone? We performed two additional sets of experiments in new mice to address this. First, we recorded neural responses (exactly as described above) in mice habituated on a locked wheel, where the potential for locomotion was completely prevented. Remarkably, dLGN responses on the locked wheel closely matched those in the tube context (Fig. S5D), with no significant differences in baseline firing rates or visual onset latencies between the two conditions (Fig. S5E, F). Second, we performed detailed analysis of postural tone (via high-speed video) in the moving wheel (stationary trials only), locked wheel, and tube; in all cases, we found that postural differences did not explain any of the findings (Fig. S6). Taken together, these results suggest that environmental context and its potential for action is a key factor that influences visual processing. These contextual differences shape the level of ongoing activity and the speed of visual responses in dLGN, both the single neuron and population levels, and emerge amidst similar arousal levels.

In summary, our study shows that environmental context itself shapes visual processing in the dorsal lateral geniculate nucleus (dLGN). These effects were revealed by examining visual neural activity in identical states of alertness and only during periods of stillness. We found elevated background activity, faster visual responses, and smaller spatial receptive fields in a context that enabled locomotion (wheel), versus one that did not (tube). Taken together, our findings reveal an unexpected influence of the physical environment on fundamental aspects of neural processing in the dLGN, with implications for how such effects of the environment may be rapidly broadcast throughout the visual system.

An important feature of our experimental design was the focus on periods of stillness across contexts, when neural responses were free from the consequences of ongoing body-wide movement. The main difference between the two contexts was that one of them (wheel) permitted locomotion. However, stationary periods were the most frequent spontaneous behavioral state on the wheel (Fig. S1J), facilitating comparison across the contexts. This natural propensity for stationary conditions is a common but often overlooked aspect of studies where mice are in a set-up that permits locomotion. In many studies, only periods of active locomotion are analyzed in order to ensure an active, engaged, low variability brain state^20–22^. In our experiments, reassuringly, we found the same locomotion-induced modulation of visual responses in dLGN on the wheel (Fig. S1B, C). However, since ongoing locomotion dramatically increases arousal level, and this affects both baseline and visually evoked firing in dLGN^9,14^, this potentially obscures the more subtle effects of the environmental context on visual processing. Similarly, eye movements are the greatest during running^8,9^, and could obscure interpretation of changes in spatial receptive field properties. These effects would be minimal during stationary trials, and we further eliminated any potential confounds of eye movements (Methods). Further, we were careful to restrict our analysis to those moments where the offset (or forthcoming onset) of running was >2 seconds away in time, since the cessation of running is followed by low-frequency thalamocortical oscillations that diminish visual responsiveness^23,24^. Thus, after carefully trying to account for these many well-known factors associated with locomotion, we were able to find that baseline visual processing was impacted by context, and that this was not simply predictable by large differences in the momentary read out of brain state, using several different electrophysiological signatures.

Regardless of our choice of arousal measure – either neural or external – we observed an overall elevation in baseline firing and faster responses in the wheel context. We primarily categorized brain state with the simultaneously recorded visual cortical delta power (1-4 Hz) and categorized trials based on the brain state in the pre-stimulus period. Cortical delta power is a major correlate of arousal mediated changes to brain state^15^ and low-frequency power co-fluctuates with changes in other arousal-related measures such as pupil size and locomotion onsets/offsets^2,6^; further, during awake behaviors, low frequency LFP power captures changes in alertness and attention even in the absence of overt movements^25–27^. Critically, we examined several other ways to measure arousal, such as narrowband gamma (NBG) power in LGN^2,28,29^, pupil size^6^, and facial motion energy^23,30^ (Fig. S3), and multidimensional analysis of multiple measures at once (Fig. S4). All these different manners of arousal categorization revealed similar effects on baseline activity and response timing as simply using cortical LFP delta power. Although we used transient stimuli and focused on response onsets, it has recently been shown that changes in arousal-related processes during wakefulness influence dLGN encoding of visual information at various timescales^31^. It is also important to note that we primarily categorized trials based on LFP from higher visual cortex, but again the main effects were consistent using other measures, including direct readouts of arousal (NBG power) in dLGN^29,32,33^. Our central finding of substantial changes in response timing (tens of milliseconds) at single neuron (Fig. 2A-F) and population levels (Fig. 2G) in different environmental contexts is broadly consistent with other studies showing millisecond-scale modulation of dLGN response timing^34^. Further, the two environmental contexts did not prevent how ongoing increases (or decreases) of arousal modulated response timing, baseline activity, or receptive field size; context mainly changed the overall set point for the relationships.

We found that in a context that permits locomotion, the visual system appears primed to process spatially localized information more quickly - even during total stillness. Further, we revealed that RF size increases as a function of increasing arousal even during stillness (Fig. 4F), echoing the effects previously observed with locomotion-induced arousal increases^9^. On the wheel, RF size was significantly smaller than on the tube; this coupled with faster latencies may allow for rapid and precise detection of high spatial frequency stimuli in an environment that enables locomotion— even during stationary epochs. Broad effects of environmental (and other) forms of context are known to affect the early visual system, and even impact feature processing at the level of the retina (e.g., spatial frequency tuning^14,35^). Our findings could imply that selectivity for visual attributes that themselves affect response latencies (e.g., spatial frequency^36^) could also differ across contexts, perhaps with further consequences for visuospatial selectivity and timing.

Several classical measures of brain state did not obviously explain the effects of the different contexts. Elevated baseline activity and faster responses on the wheel suggest that perhaps an overall DC depolarization of dLGN neurons drives the effects. Effects of DC depolarization may not be readily apparent in frequency-resolved electrophysiological measures within dLGN, nor in the cortical LFP readouts of brain state; likewise, they may not be apparent in slower external behavioral indicators of arousal such as pupil or facial motion. What could potentially drive this depolarization? A first possibility is cholinergic input, either directly or indirectly, through activation of the basal forebrain (BF). Indeed, tonic cholinergic stimulation slowly depolarizes dLGN neurons^37^, and activation of BF increases baseline activity in dLGN and improves trial-by-trial visual response timing, enhancing reliability^38^; it is conceivable that cholinergic activity contributes to the different contextual effects on baseline activity and response timing in our experiments. An experiment to test this would be to record from dLGN in mice in the tube, then apply cholinergic agonists in dLGN locally (or by activation of the BF) and see if this mimics increased baseline activity and faster response latencies as found on the wheel. However, in monkey V1, acetylcholine shrinks RF size^39^, unlike arousal mediated changes in mouse dLGN and in V1^9,40^, possibly due to non-monotonic effects on excitability^6^. A second source of depolarization could be long-lasting activation of the mesencephalic locomotor region (MLR) that persists during stationary epochs on the wheel; for example, subtle activation of the parabrachial region within the MLR (that does not elicit locomotion) nonetheless reduces dLGN burst firing^4,41^, as we observed here. A third potential source of depolarization is the noradenergic system; in V1, this maintains tonic depolarization during locomotion^24^; likewise, noradrenergic agonists in dLGN reduce burst firing^42–44^. Lastly, depolarization of dLGN neurons could arise from V1 feedback^19,45^. However, V1 provides both direct excitation and indirect inhibition of dLGN via the thalamic reticular nucleus, so this may not result in a straightforward, DC depolarization of dLGN^46,47^. Beyond the observed effects here of higher baseline activity and faster response timing, a context-mediated depolarization could alter other aspects of visual processing. First, as argued above, depolarization could alter the degree of burst firing in different contexts^7,9,31^; indeed, we found burst firing was reduced in the wheel versus tube (Fig. S2A,B). Second, baseline depolarization could expand the dynamic range available to encode luminance changes^48,49^. This regime may be advantageous for dLGN neurons to signal rapid changes in luminance in contexts that enable interleaved locomotion and stillness. Future experiments across contexts could disentangle these and other aspects of thalamic contextual modulation.

There are several limitations to our study which readily suggest future experiments in different contexts. One main limitation of our experimental design was the inability to record from the same neurons across each context. This concern is somewhat mitigated by the fact that, in the majority of our experiments, recordings in each context were performed within the same mice within days of each other, and within animal, the effects of context were consistent (Fig. S7, Table S1). Future experiments utilizing chronic extracellular recordings in dLGN or imaging of dLGN activity would enable the comparison of the same neurons in one context versus another. A second limitation is that the contextual effects described here could be explained by factors that were not observable with electrophysiology. For example, locomotion can induce changes in body temperature, which influences neural excitability and synaptic function at a longer timescale beyond the motor event^50^. Slower-acting intracellular signaling cascades such as those involving neurotrophins may result in longer-lasting effects on neural responsiveness^51,52^. Future work could assess such contributions to the context-dependent shifts in baseline activity and response latency in dLGN. A third limitation is our choice of stimulus set. While sparse noise stimuli are excellent for assessing changes in receptive field properties, a constraint is the limited number of trials in which the best stimulus appears in a given neuron’s RF. This hinders assessing the variability of RF responses across multiple timescales. In future experiments, tailored stimuli (i.e., repeated, frequent stimuli in the RF alone) would allow for more detailed investigation of how arousal influences dLGN activity over multiple temporal scales^31^.

In summary, we revealed that the overall physical environment shapes basic features of visual processing in the thalamus. Contextual effects are typically thought to be strongest in cortex^53–56^ so why might contextual modulation of LGN convey ecological advantages? In freely behaving mice, stationary epochs are naturally punctuated by running behaviors, such as during prey-capture and escape^57,58^. The visual system may thus be tuned to process information more quickly in contexts that promote action — even when mice are totally still — so that ensuing visual processing during locomotion is already primed for integration of higher temporal frequencies, optic flow, etc. It would be most beneficial to accelerate the timing of visual signals at the earliest stage, rather than individually in multiple downstream structures. This quickened regime on the wheel is not readily observable in experimental contexts (tube, locked wheel) that nonetheless survey the system in comparable motor and arousal states. However, by limiting the potential for locomotion, the tube context also minimizes widespread neural effects that may pose other interpretational complications for the experimental question at hand, such as in psychometric studies of stimulus detection^3,8,59,60^. Overall, our findings highlight that the experimental choice of physical environmental context can have large impacts on the most basic aspects of visual processing, even as early as the primary visual thalamus. The information about environmental context sculpts baseline activity, response timing, and spatial information, allowing for more integrative processing of contextually relevant visual information to be broadcast to the rest of the visual system. More broadly, our findings imply that inherent properties of the environmental context could have marked effects on setting the neural information processing regime across other sensory systems.

## Acknowledgements

We thank Tony Lien and Elaida Dimwamwa for feedback, David Weiss and Garrett Stanley for sharing materials, and Tiernon Riesenmy and Jeff Markowitz for assistance with pose tracking. This work was supported by National Eye Institute (F31EY033691 to K.P.), National Institute of Neurological Disorders and Stroke (T32NS096050 to K.P.; R01NS109978, RF1NS132288 to B.H.)

## Author Contributions

K.P. performed silicon probe recordings, wrote code and analyzed data; K.P., B.H. wrote the manuscript.

## Declaration of interests

The authors declare no competing interests.

### Lead contact

Further information and requests for resources should be directed to and will be fulfilled by the lead contact, Bilal Haider.

### Materials availability

This study did not generate new unique reagents

### Data and code availability

All data structures and code that generated each figure will be publicly available at DOI: 10.6084/m9.figshare.26364601 upon publication and linked from the corresponding author’s institutional webpage upon publication.

## STAR Methods

### Experimental subject details

| Reagent or Resource | Source | Identifier | Number of Mice | Number of Recordings [wheel;tube] |
| --- | --- | --- | --- | --- |
| **Experimental Models: Organisms/Strains** |  |  |  |  |
| **Mouse C57BL/6J** | **Jackson Laboratory** | IMSR_JAX:000664 | 8 | [7;17] |
| **Mouse Ai32 x Scnn1a-Cre** | **Jackson Laboratory** | IMSR_JAX:009613 | 1 | [0;1] |
| **Mouse B6.129S4-Gt(ROSA)26Sortm1Sor/J** | **Jackson Laboratory** | IMSR_JAX:003474 | 2 | [6;2] |

All experimental procedures were approved by the Institutional Animal Care and Use Committee (IACUC) at the Georgia Institute of Technology. Three male mouse genotypes were used: C57BL/6J, Ai32 x Scnn1a-Cre, and B6.Gt(ROSA). Mice were housed with littermates until experiments, then single-housed in enriched environments (nesting material, tubes, and blocks). Selection for experiments was random, based on age and health, and housing followed a 12h-12h light-dark cycle.

### Method Details

#### Implant surgery

Mice were implanted with a custom-built stainless steel head plate containing a recording chamber (3 mm in diameter) during isoflurane anesthesia (3% induction, 1-2% maintenance). The head plate was fixed to the skull using a layer of veterinary adhesive (VetBond) before being secured to the cranium with dental cement (Metabond). The recording chamber was sealed using an elastomer (KwikCast). Following implantation, mice were given a minimum of 3 days to fully recover.

#### Habituation to recording apparatus

Mice in the tube context only (4/9) were gently handled and habituated to the recording environment for a minimum of three days prior to experimentation. For mice that were habituated in both contexts (5/9), we randomized the starting context (tube first or wheel first) in order to minimize any systematic biases. Total habituation to both contexts took between 6-8 days. Regardless of context, mice were slowly acclimated to the environment. On the first day, mice were habituated for 30-45 minutes. On the second day, 60-90 minutes, and on the third day, 120-180 minutes. Starting on the second day and through the rest of the habituation period, mice were exposed to visual stimuli (small flashed squares). Animals were considered to be fully habituated when they were able to stay on the recording apparatus for a full 180 minutes (typical experiment length) with no signs of agitation (elevated tail, fluid buildup in eye, vocalizations). Habituation followed the same procedures for locked wheel experiments, where animals were never exposed to the wheel moving during habituation.

#### Stimulus Display

Visual stimuli were displayed on gamma-corrected LCD displays (Dell Ultrasharp U2417H or U2419H, 60 Hz refresh rate) with a peak luminance of 250 cd * m^−2^. Two displays were positioned at right angles to one another such that stimuli at 0° and 90° in azimuthal space have similar viewing angles relative to the mouse’s eye. To confirm linearization, we displayed full contrast stimuli (full black to full white) and measured the resulting monitor luminance values with a photodiode (Thorlabs), then corrected this relationship with the (inverse) exponential function. We measured light levels using a photometer (AEMC CA811) with spectral sensitivity range (500 – 620 nm) overlapping the peak absorption wavelengths for both rods and M-cones, positioned at the same viewing angle as the mice. Measurements from the experimental apparatus were averaged (±SD); stimuli at [100% black, 50% gray background, 100% white] stimuli provided [0, 117.5 ± 14.4, 237.3 ± 28.7] cd * sr * m^−2^.

#### Visual stimuli: Full Screen Flash

Stimuli were created using Matlab (2017b, 2019a) with the Psychophysics toolbox. Full screen flashes spanned from approximately − 60° to + 150° in azimuth. Stimuli consisted of a background of mean luminance (gray) for 0.25 s followed by either black (minimum luminance) or white (maximum luminance) screen for 0.25 s. Each full screen flash session consisted of 500-1000 trials (Fig. S1).

#### Visual stimuli: Receptive Field Mapping

Stimuli were created using Matlab (2017b, 2019a) with the Psychophysics toolbox. Receptive field (RF) mapping stimuli consisted of individually presented black (minimum luminance) or white (maximum luminance) against a gray background of mean luminance. Each square was 7°x7° and appeared in one of 160 spatial locations covering 90° (azimuth) x 50° (elevation), spanning the monocular and binocular visual field. Each stimulus was presented randomly across the 160 given stimulus locations, such that the same square would not appear in the same location on the subsequent trial. Stimuli were displayed for 0.1 s followed by a 0.3 s inter-stimulus interval. Stimuli were presented in blocks of trials, such that a block consisted of a single luminance of the square, and each square was presented in each location 10 times within one block. Each experiment consisted of 10-30 repeats of square stimuli of either black or white in each spatial location.

#### Recordings: dLGN and visual cortex

dLGN was targeted with stereotaxic coordinates (−2.5 mm posterior to bregma; −2.5 mm lateral from midline^61^. Each Neuropixels 1.0 recording electrode contains a total of 960 channels, of which a subset of 383 were used for recording. Spikes were acquired at 30 kHz and local field potential (LFP) at 2.5 kHz via a PXIe Card, a National Instruments board, and Spike GLX software. In all experiments, a full field flash stimulus was shown while advancing to dLGN (2.8 - 3.2 mm below the dura) to confirm visual responses, and in most instances confirmed with histology. We note that the time of day neural recordings took place did not differ across experimental contexts (p = 0.959, Kolmogrov-Smirnov test)^52^.

#### Recordings: Animal position

The head fixation apparatus in the recording environment was identical across all experiments. Mice in each context were always placed in the same position to ensure that viewing distance from the stimulus screens were identical across experiments (see Fig. S6).

#### Recordings: Eye camera

Face and eye movements were monitored by illuminating the animal’s face and eye with infrared light (Mightex, SLS-02008-A). The right eye was monitored with a camera (Imaging source DMK 21Bu04.H) fitted with a zoom lens (Navitar 7000) and a long-pass filter (Mightex, 092/52×0.75). The camera was placed ~22 cm from the animal’s right eye. Videos were obtained at 30 Hz.

#### Recordings: Locomotion

In wheel experiments, mice were free to run on a circular treadmill. The circular treadmill contained four equally spaced dowels around the perimeter. Running events were measured by an optical detector that triggers events following an IR beam break. As the wheel rotated, the dowels caused rapid beam breaks and triggered pulses. Running speed was quantified as the circumference of the wheel divided by the rate of trigger pulses per second. Trials were considered as locomoting if the calculated speed exceeded 1 cm/s for ≥ 1 s; trials were considered stationary when the speed was 0 cm/s for ≥ 1 s. Based on these metrics, each trial was categorized as: 1) stationary, 2) locomotion, 3) within 1 s of locomotion onset, 4) within 1 s of locomotion offset. Trials that did not meet these criteria (running onsets/offsets; trial types 3 and 4) were excluded from analysis (11% of trials; Fig S1J). The total fraction of time spent running per session (within mouse) did not explain any of the main findings during stationary trials (correlation between fraction of time running in session versus baseline activity: r = −0.205, p = 0.733; between fraction of time running in session versus onset latency r =0.600, p = 0.350).

#### Recordings: Atropine

For recordings during atropine application (Fig. S2F, G), 1-3 drops of 1% Atropine Sulfate solution (Alcon) were applied topically on the eye. Recordings were performed after the pupil was fully and stably dilated, ~30 minutes after application (Fig. S2F). Full screen flash stimuli (both black and white) were presented to ensure the lack of light-induced pupillary responses.

#### Histology

In some experiments, prior to LGN recordings, lipophilic dye (DiI, Invitrogen) was painted on the back side of the recording electrode and inserted in dLGN for a minimum of 2.5 hours. At the end of the recording, mice were transcardially perfused with 4% paraformaldehyde (PFA, VWR) in 1x PBS. Brains were kept in 4% PFA in 1x PBS for 24 hours and transferred to PBS. Brains were sectioned on a Vibratome (Leica), and each section was 100 um in thickness. Brains were then stained with DAPI (2 mM in PBS, AppliChem). We then mounted each slide with a fluorescent mounting medium (Vectashield, Vector Labs). Sections were imaged using a Confocal microscope (Zeiss). Targeting of dLGN was confirmed post-mortem, and any data obtained from mice where we observed a recording track outside of dLGN was discarded.

### Quantification and Statistical Analysis

All data analysis was performed in MATLAB. Statistical significance was set at (α) < 0.05 for all comparisons.

#### Data analysis: Spike sorting

Single units in dLGN were isolated and identified using Kilosort2^62^. Manual curation of clusters was performed using Phy2^63^. Clusters were chosen if the refractory period violations (occurring within 2 ms) occurred in fewer than 1% of all spikes assigned to a cluster^64^.

#### Data analysis: Local Field Potential

A single-shank Neuropixels 1.0 electrode was used to simultaneously record the local field potential (LFP) in visual cortex and single unit activity in dLGN (Fig. 1C). Here, visual cortex was either the binocular portion of primary visual cortex (V1), rostrolateral (RL), or anterolateral (AL)^65^ higher visual cortex. Visual responses to full screen flash stimuli were confirmed by inspecting the real-time neural data streaming during recordings, but the exact area boundaries and retinotopy of recorded regions within the visual cortical areas were not ascertained. We used LFP delta power in the visual cortex as a measure of changes in arousal. We used the spectrogram function in Matlab (Mathworks) and defined delta power as the power in the 1 to 4 Hz frequency band. We calculated the power over a sliding 1 s window with a 90% overlap to achieve 0.1 s precision. Next, we normalized the power spectrum by the sum across all frequencies at each time point, giving the relative power in the delta band^19^. We then smoothed the resulting delta power vector over a 1 s moving window. Pre-stimulus delta power was then calculated for each trial.

#### Data analysis: Arousal Level

In order to control for arousal level, the pre-stimulus delta power distribution was taken where the data had the greatest overlap between the tube and wheel context, which occurred at a moderate level of arousal (0.03 – 0.08 normalized delta power; full range: 0-0.15 normalized delta power) (Fig 1E).

Pre-stimulus delta power was calculated across all recordings for both contexts and plotted as a histogram (Fig. 3A). The histogram was then split into five equally sized bins, denoting the five increasing levels of arousal. When defining arousal levels within each context, the same bin widths and delta power values were used to define each level of arousal.

To make our analysis more stringent and exclude sampling differences between distributions within each quintile, we tried to match the distributions by applying mean-matching^66^. Unfortunately, this procedure required dropping 30.3% of trials total in the wheel context, where we have the fewest data points. As a result, the data set became under-sampled and too limited to draw interpretable conclusions.

#### Data analysis: Pupil and periocular movements

To fit pupil size and calculate eye position, we used DeepLabCut^67^. Pupil size was tracked using eight points around the circumference of the pupil. Motion energy was calculated as the sum total of the absolute difference in pixel intensity between adjacent frames^30^. 8 points were fit around the eye from the bottom to top lid and the inner and outer corners. These pixels were excluded so that changes in pupil size do not contribute to the facial motion energy vector. At the end of every experimental session, mice were given liquid condensed milk while viewing a gray screen. Liquid consumption was chosen as a calibration event, because it evokes a large and standardized amount of motion energy and pupil dilation that is consistent between tube versus wheel experiments. In wheel experiments, we ensured mice were not actively running prior to consuming the milk, and during consumption animals were not actively running. We used the maximally evoked pupil area during consumption to normalize all pupil sizes as a fraction of the maximum evoked by reward consumption. We also normalized facial motion energy values to the same event, and normalized all motion energy values to the maximum motion energy value during consumption.

#### Data analysis: Permutation test

To ensure that our analysis of timing was statistically robust and representative of the underlying activity across recordings and not driven by individual animals, recording days, or units, we analyzed resampled populations of neural recordings. Surrogate populations (Fig 2G) were created by first randomly sub-sampling all units across recordings (with replacement, 40% of units). This procedure was performed the same number of times as the number of recordings in each context (17 datasets for tube, 6 datasets for wheel). We averaged the surrogate datasets to create a population RF map over time. The activity from the center pixel of the population RF map was selected and turned into a peri-stimulus time histogram. Test statistics were calculated for onset latency, peak latency, and response duration between tube and wheel. 500 permutations were performed. A p value was calculated by counting the number of test statistics greater than the initial test statistic, divided by 500. A result was significant at p < 0.05.

#### Data analysis: ON/OFF classification in dLGN

Neurons in dLGN were classified as having ON or OFF preference from the response to black or white squares flashed in the neuron’s receptive field. The number of spikes evoked by the change in luminance (40 – 140 ms after stimulus onset) were counted and used to calculate the ON/OFF index.

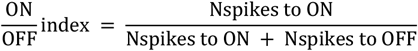

Neurons with an ON/OFF index of < 0.3 were identified as OFF cells and > 0.7 were ON dominant cells, as in previous studies^29,68^; wheel: 17 ON, 15 OFF, 40 mixed selectivity; tube: 72 ON, 48 OFF, 115 mixed).

#### Data analysis: dLGN Receptive Field Analysis

First, grand average receptive field (RF) maps were created for each neuron. RF maps were created by generating a 2D histogram of spike counts for every neuron at each of the 160 spatial locations of the mapping stimulus. Maps were binned at 0.01 s. Both black and white responses were combined to create grand average RF maps. The average response between 0.04 – 0.14 s after stimulus onset was taken for every unit to create a receptive field map^69^. A Chi-squared test for independence was used to determine whether a map had locations with significant (p < 0.05) responses above baseline that together defined the receptive field^56^. Except for Fig. S1 (all visually responsive dLGN neurons), only those neurons with a significant spatial RF map were analyzed in this report (49% of all recorded neurons with visual responses to full screen flashes also passed significance criteria for 2-D RF maps, consistent with prior work^56^). We note that the proportion of significant RF maps was highly similar across conditions: 49% in tube (276/585), 49% in wheel (107/218, p = 0.644, Chi square). Most neurons had RF elevation locations near eye level (0°) in both contexts (tube: 8° ± 13°; wheel: 8° ± 13°, p = 0.930). Azimuth locations differed, with neurons in the tube biased towards binocular visual space (tube: 41° ± 25°; wheel: 56° ± 29°, p = 2.75e-5). After controlling for this bias by examining only monocular neurons (tube: mean RF location 68° ± 15°; wheel: mean RF location 75° ± 13°; p = 0.061), the significant differences in our main findings on baseline activity (p = 0.002) and onset latency (p = 0.001) persisted between the two contexts.

#### Data analysis: Arousal-based Receptive Field Maps

To create arousal-based RF maps, the grand average RF maps were separated by trial type. Each trial was assigned a pre-stimulus delta power value over the 1 second prior to stimulus onset within the neuron’s receptive field. The distribution of delta power over all experiments was taken and separated into three equally sized bins to determine ‘low’, ‘mid’, and ‘high’ levels of delta power. Trials that fell into each category were then taken to create the arousal-based maps for each neuron (Fig. 4 F, G; example maps generated from matched arousal trials as in Figs 1,2).

#### Data analysis: Receptive Field Center Size

To calculate receptive field center size, individual receptive field maps were first baseline subtracted (0.1 s pre-stimulus activity) and the center of the receptive field was identified as the spatial location where the most spikes were evoked to the presentation of the sparse noise visual stimulus. Any pixels contiguous with the center pixel that evoked spikes at greater than or equal to 50% of the maximally evoked center pixel were included as part of the receptive field. Area was calculated assuming RF circularity (A = π r^2^). Eye movements still occur in stationary animals, although to a lesser degree^70,71^. Therefore, we also restricted our receptive field analysis to trials where the eye position deviated less than the size of the visual stimulus (within 7° of the average central position across the recording, as in^70^). This excluded just 4.3% of all trials on the tube, and 2% of all stationary trials on the wheel.

#### Data analysis: Spatial Selectivity

To calculate receptive field spatial selectivity, we used a modulation index (MI) to assess firing at the center of the receptive field relative to the background for each individual neuron. Using the same ‘center’ pixels as identified for the receptive field, the average firing rate in the receptive field was calculated for all of these, while all other pixels (typically 155-159 outside of the center; from 160 total pixels in a single RF map) were taken as background. Here, we did not baseline-subtract the receptive field maps in order to capture the differences in the actual firing rates within neuron.

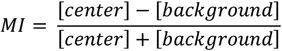

#### Data analysis: Quantifying visual response amplitude and timing

All peri-stimulus time histograms (PSTH) had a bin size of 0.01 s. Due to inherent variability and sampling constraints, neurons with significant RF maps could nonetheless fail to have a stimulus appear in the receptive field during a particular condition (e.g., stationary trials within one of the five binned levels of arousal). To calculate baseline firing (Fig 2B, Fig 3 D-F), we took all neurons that had a significant receptive field map (n=276 tube, n=107 wheel). For all other stimulus-evoked analysis, the response to the best stimulus location and preferred color (black or white) was taken for each neuron. When analyzing baseline activity for those subsets of neurons that also had a response to the best stimulus location within a particular condition, we still observed a significant effect of context on baseline activity (data not shown). If a neuron was not statistically categorized as ON or OFF (i.e., mixed selectivity), average responses to both black and white stimuli were analyzed. Baseline activity (Fig. 2B; Fig 3D E, F) was calculated in the 0.1 s prior to stimulus onset. Baseline-subtracted peak visual response (Fig. 2C; Fig. 3G-I) was calculated by subtracting the baseline activity (0.1 s prior to stimulus onset) from the average firing rate in a 0.05 s window 0.04 s after stimulus onset. The onset latency (Fig. 2D; Fig. 3J) was calculated as the earliest time after stimulus onset during which baseline-subtracted cumulative PSTH was above a 95% bootstrapped confidence interval on the cumulative baseline values^72^. The peak response latency (Fig. 2E, Fig. 3K) was the time at the maximum evoked response for each individual neuron. The response duration (Fig. 2F, Fig. 3L) was quantified by assessing the duration of the interval in which the response was at or above 25% of the maximum firing rate. A few neurons with response durations >0.2 s were excluded from analysis (1/72 units on wheel, 3/235 units on tube).

#### Data analysis: Bursting

Bursts were defined as 100 ms of silence followed by >= 2 spikes within 4ms^9,73^. The burst ratio was then defined as the number of burst spikes/total spikes.

#### Data analysis: Postures

Neural recordings were conducted simultaneously with high-speed full-body videography (M3Z1228C-MP 2/3” 12-36mm F2.8 Manual Iris C-Mount Lens, Computar, 30fps, Fig S6). Body keypoints, including left and right hands, nose, and the base of the tail, were labeled using the SLEAP deep-learning body tracking algorithm^74^, and tracked the trajectories of these keypoints across all frames. The motion capture camera and head-fixation apparatus were identically securely mounted to the air table in both contexts to ensure consistency across recordings. During electrophysiological recordings, we recorded LFP, pupil, and dLGN spiking activity while presenting the same sparse noise stimuli described in “Visual Stimuli: Receptive Field Mapping.” Stationary trials were identified in the wheel context (as described in “Recordings: Locomotion”), and we selected corresponding trials in both wheel and tube context that overlapped in pre-stimulus delta power distribution as in Fig. 1E. Trials were then sorted by the amount of keypoint displacement (D_*i*_), defined as the total combined (*x, y*) displacement of the left hand (LH), right hand (RH) and rear quarter (Rear) keypoints on each frame (*i*), relative to the mean 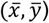 positions across all frames.

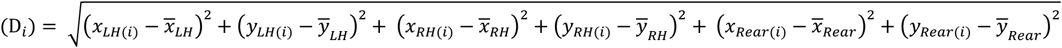

Non-overlapping displacements (suggesting distinct postures) were defined as those where less than 1% of total trials of one context overlapped with the other one (Fig. S6; tails of distributions). The remaining (central) portion of the distributions were classified as overlapping displacement (suggesting similar postural tone).

---

**Supplementary Figure 1:**
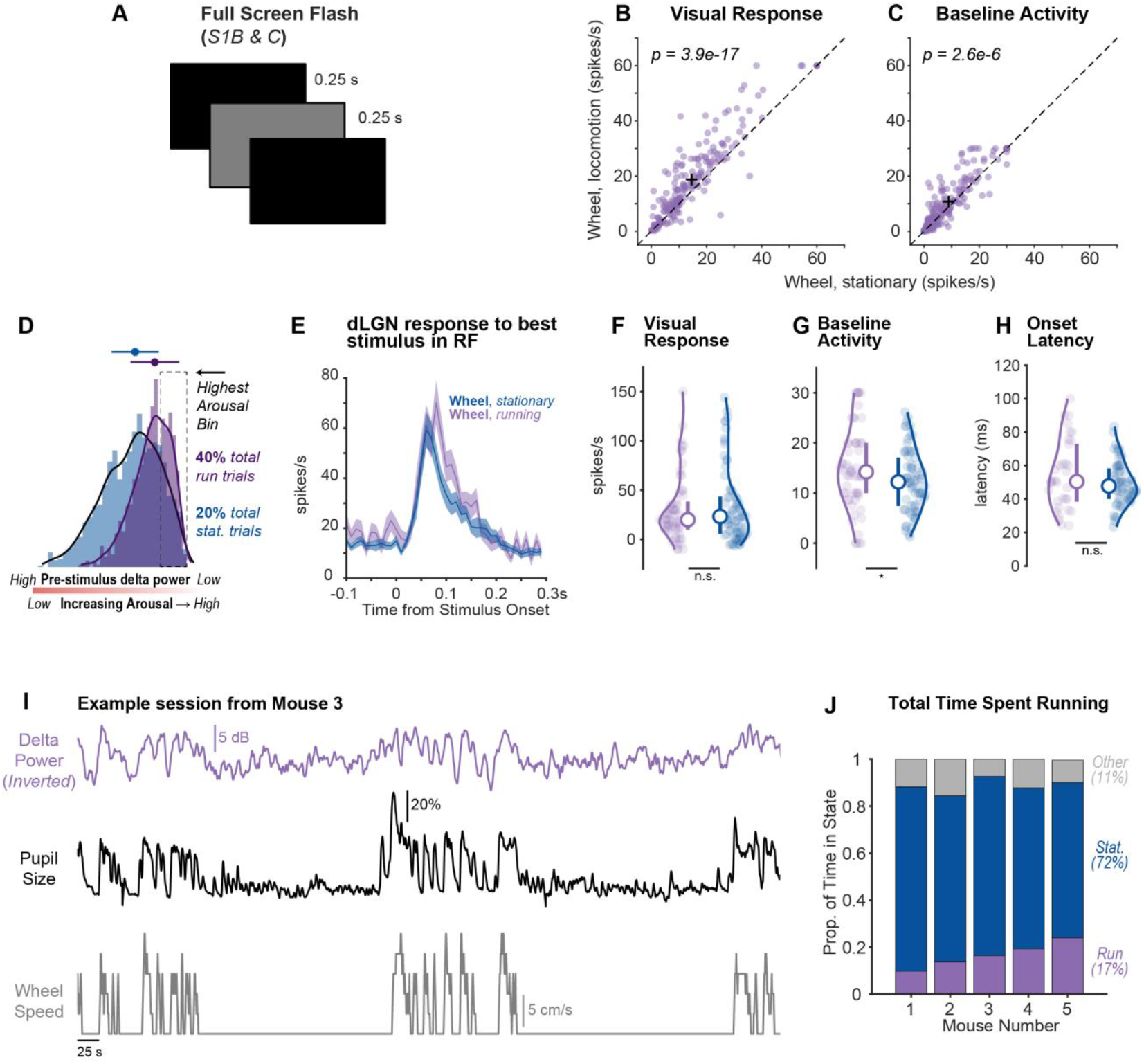
Visual responses on wheel during running versus stationary trials. **A**. Full screen flash stimulus (grey to black illustrated here; grey to white also shown in experiments). **B**. Visual response of dLGN neurons to a full screen flash stimulus is elevated during locomotion versus stationary epochs (N = 5 mice, n = 196 neurons, *p* = 3.9e-17; sign rank test). **C**. Baseline activity of neurons during gray screen periods is elevated during locomotion versus stationary epochs (*p* = 2.6e-6; sign rank test). **D**. Distribution of delta power preceding all analyzed visual stimulus trials on the wheel for stationary trials (dark blue; 2386 trials) and running trials (purple; 1274 trials). Note that delta power axis is reversed (arranged high to low), so that arousal level increases from left to right. 511/1274 (40%) of trials during running occur within the high arousal bin (dashed box, see Fig. 3A). Conversely, 465/2386 (20%) of stationary trials occur within the high arousal bin. **E**. Average dLGN spiking response on wheel during running (purple) vs. stationary (dark blue) trials when the best stimulus was presented in a given neuron’s receptive field. **F**.Visual response amplitude is similar between running and stationary trials (running: 20 ± 14 spikes/s, n = 64; stationary, 23 ± 18.5 spikes/s, n = 72; Median ± IQR/2; *p =* 0.494; Wilcoxon one-sided rank sum test throughout figure). **G**.Baseline activity greater during running versus stationary trials (run 14.2 ± 5.0 spikes/s, stationary 12.22 ± 5.2 spikes/s; *p* = 0.033). **H**.Onset latency was comparable between running and stationary trials (*p* = 0.204). **I**.Example session over a single recording block (~12 minutes for a full block of squares tiling the visual hemifield). Traces show aligned visual cortical LFP delta power (inverted, purple), pupil size (black), and running wheel speed (gray). **J**.Proportion of time spent running on average across all recording sessions for each given mouse recorded in the wheel. Across all recording days and mice, mice on average ran 17% of the time, and were stationary 72% of the time. On average, 11% of trials were excluded as uncategorized for either running or stationary criteria due to being within 1 s of a running onset/offset (see Methods).

**Supplementary Figure 2:**
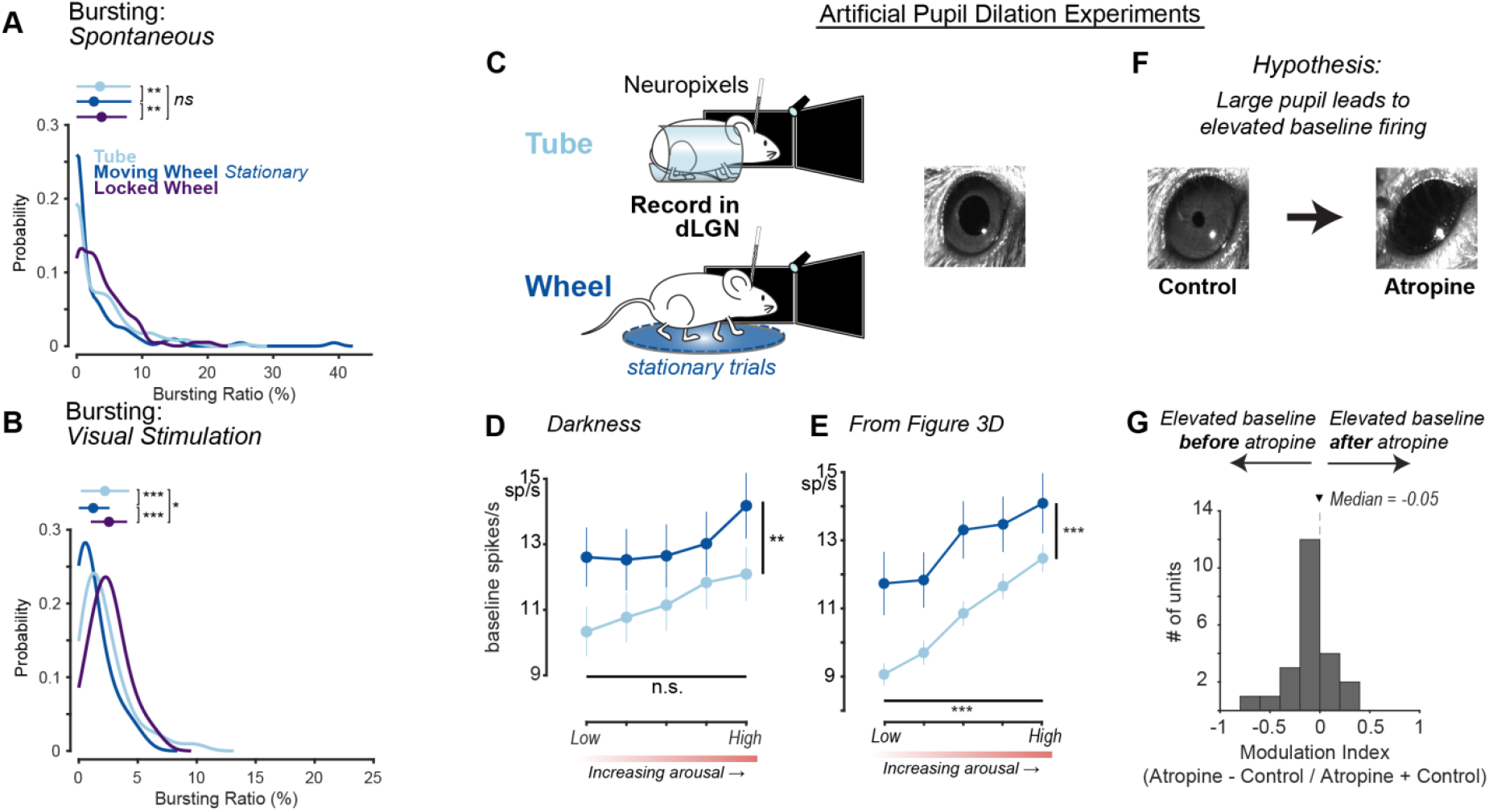
Analysis of bursting, and artificial dilation of pupil does not increase spontaneous firing rates. **A**. The spontaneous bursting ratio is significantly modulated by context (p = 0.00014, Kruskal-Wallis ANOVA). Bursts defined as 100 ms of silence followed by >= 2 spikes within 4ms, ratio defined as number of burst spikes/total spikes. Post-hoc tests reveal burst ratio is not significantly different between tube and locked wheel (tube: 3.6 ± 4.5%, n = 235 neurons; locked wheel: 3.8 ± 3.8%, n = 86 neurons, p = 0.116, Bonferroni tests) but it is significantly reduced in the moving wheel, both relative to tube (tube: 3.6 ± 4.5%; moving wheel 2.6 ± 5.7%, n = 72 neurons, p = 0.007) and relative to locked wheel (locked wheel: 3.8 ± 3.8%; moving wheel 2.6 ± 5.7%, p = 9.45e-05). **B**. Visually-driven bursting (best stimulus in RF for a given neuron) is significantly modulated by context (p = 2.33e-10, Kruskal-Wallis ANOVA). Post-hoc tests reveal it is significantly reduced on tube relative to locked wheel (tube: 2.2 ± 2.0%, n = 235 neurons; locked wheel 2.6 ± 1.5%, n = 86 neurons, p = 0.01, Bonferroni tests), and also significantly reduced on moving wheel, both relative to tube (moving wheel 1.2 ± 1.4%, n = 72 neurons; tube: 2.2 ± 2.0%, p = 9.93e-07) as wheel as to locked wheel (moving wheel:1.2 ± 1.4%; locked wheel: 2.6 ± 1.5%, p = 2.44e-10). **C**.Experiments were performed on the tube and the wheel in full darkness to elicit maximal pupil dilation on all trials (N = 2 mice, tube context; N = 2 mice, wheel context). **D**.Baseline activity on all stationary trials remained significantly elevated on the wheel relative to the tube (wheel n = 102 neurons, tube n = 90 neurons, *p* = 0.0002). There was no significant effect of increasing arousal on baseline activity in full darkness. **E**.Data from Figure 3D when animals were in front of a gray screen. **F**.Atropine was applied externally to mouse’s eye to artificially dilate the pupil (n = 1 mouse, 23 neurons). If increased baseline firing is solely due to more light entering the eye with a large pupil, atropine should increase baseline firing relative to control. Note extreme dilation of pupil after atropine relative to baseline, and compared to full darkness (in A). **G**.Baseline firing rates were not elevated when pupil was artificially dilated with atropine, relative to control firing rates in the same neurons before atropine application. Modulation expected to be positive if atropine increased baseline firing rates.

**Supplementary Figure 3:**
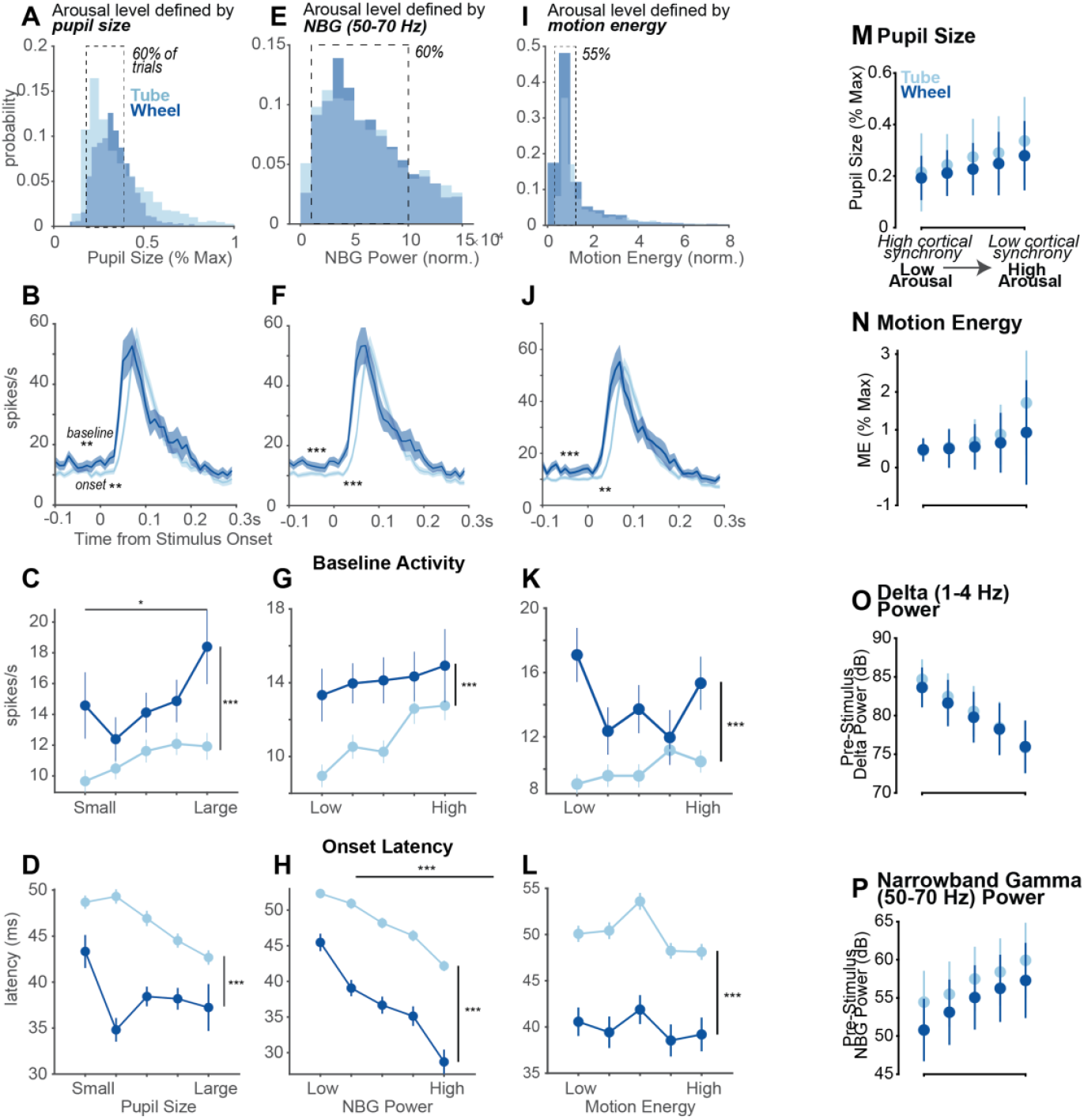
Alternative methods to define arousal level using pupil size, narrowband gamma power (NBG), and motion energy. **A**. Distribution of pre-stimulus pupil size across the tube (light blue) and wheel (dark blue) physical contexts. The center 60% of trials were analyzed. **B**. Population peri-stimulus time histogram (PSTH) of neurons in dLGN when a given neuron’s preferred stimulus is presented in the best location. Baseline activity was significantly elevated on the wheel relative to the tube (wheel n = 73, 12.9 ± 4.2 spikes/s; tube n = 219, 9.2 ± 5 spikes/s; *p* = 0.0016; Median ± IQR; Wilcoxon rank sum test; same neuron numbers and statistics throughout unless otherwise noted). Onset latency was faster in the wheel context (wheel 48 ± 7.5 ms; tube 56 ± 14 ms; *p =* 0.0021). **C**. Baseline activity was overall elevated on the wheel relative to the tube (*p* = 8.8e-6) and increased with increasing arousal (*p* = 0.0303; two-way ANOVA). **D**.Onset latency was overall faster in the wheel relative to the tube (*p* = 8e-10, two-way ANOVA). There was no significant effect of increasing arousal level on onset latency (*p* = 0.0725). **E**.Same as in A, but for pre-stimulus narrowband gamma (50-70 Hz) power. The center 60% of trials were analyzed. **F**.Same as in B. Baseline activity was significantly elevated on the wheel relative to the tube (wheel 12.0 ± 4.4 spikes/s, tube 8.8 ± 3.9 spikes/s; *p* = 7.2e-4) **G**.Same as in C. Baseline activity was significantly elevated on the wheel relative to the tube (*p* = 8.7e-5). There was no significant effect of increasing arousal level on baseline activity (*p* = 0.1339; two-way ANOVA). **H**.Onset latency was overall faster in the wheel relative to the tube (*p* = 5.5e-21, two-way ANOVA). Increasing arousal level resulted in faster onset latencies in both contexts (*p* = 6.2e-10). **I**.Same as in A, E, but for pre-stimulus motion energy (Methods). The center 55% of trials were analyzed. **J**.Same as in B, F. Baseline activity was significantly elevated on the wheel relative to the tube (wheel 13.92 ± 9.0 spikes/s, 10.1 ± 7.1 spikes/s; *p =* 5.2e-4). Onset latency was faster in the wheel context (wheel 48 ± 7.5 ms; tube 56 ± 14 ms; *p* = 0.0078) **K**.Baseline activity was elevated in the wheel relative to the tube (*p* = 2.0e-8) but was not impacted by increasing arousal level (*p* = 0.2164; two-way ANOVA). **L**.Onset latency was overall faster in the wheel relative to the tube (*p* = 1.56e-14, two-way ANOVA). There was no significant effect of increasing arousal level on onset latency (*p* = 0.1799). **M**.Pupil size across the exact same trials in the five defined arousal bins (in Figure 3A). **N**. Same as in M, but for motion energy. **O**.Same as in M, but for Delta power (1-4 Hz). **P**.Same as in M, but for narrowband Gamma power (50 – 70 Hz).

**Supplementary Figure 4.**
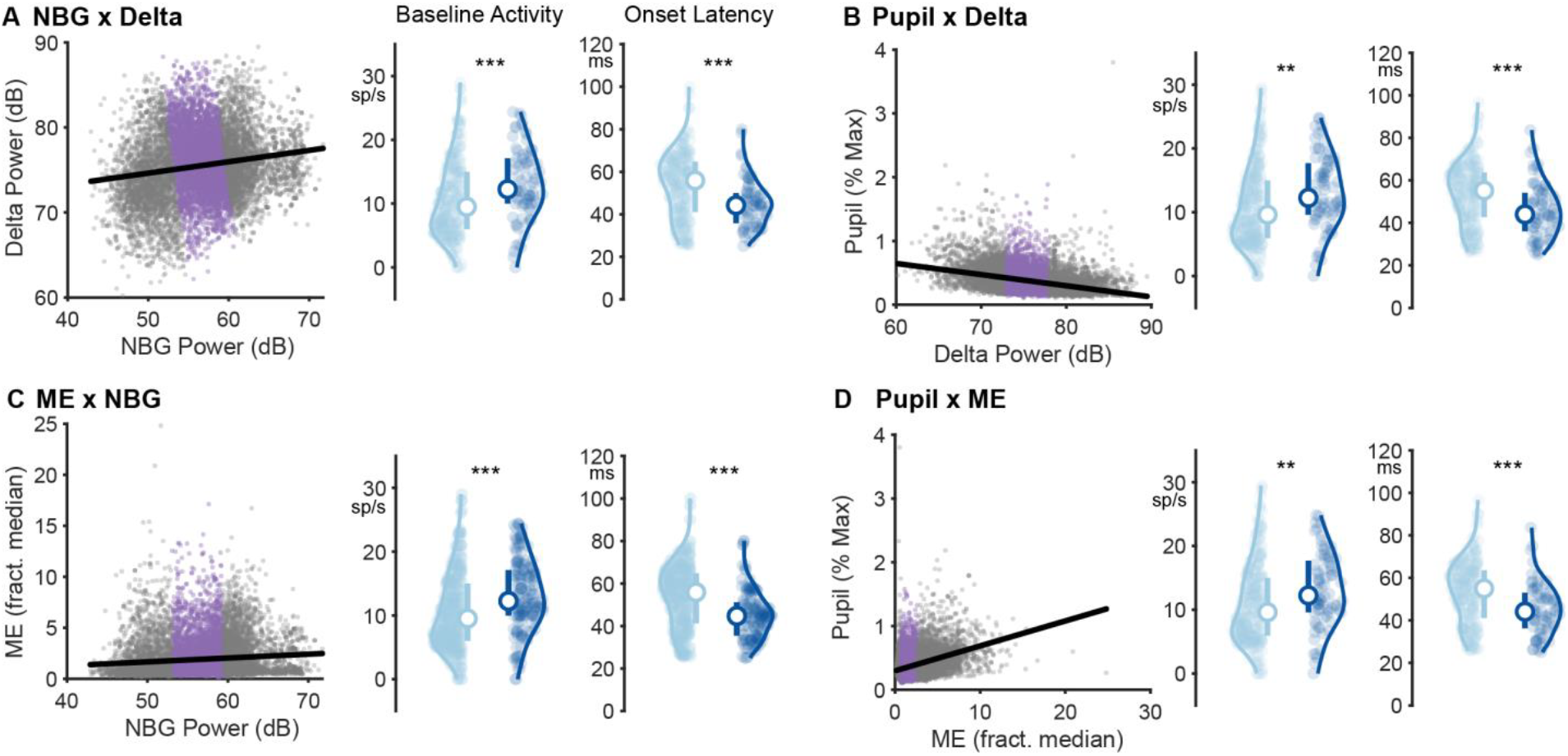
Multidimensional analysis shows same trends as delta power classification. **A**. Two-dimensional scatter plot of single trial cortical LFP narrowband gamma (NBG; 50 – 70 Hz) versus delta (1 – 4 Hz) power, for all experiments in tube (light blue) and moving wheel (stationary only; dark blue). Purple shows the central 50% of the two-dimensional distribution. Using these overlapping trials, baseline activity was elevated in the wheel (wheel: 12.3 ± 3.6 spikes/s, tube: 9.5 ± 4.5 spikes/s, n = 235 units; n = 72; p = 7.25e-5, Median ± IQR/2, Wilcoxon one-sided rank sum test throughout figure), and onset latency was significantly faster (wheel: 44 ± 7 ms, tube: 56 ± 12 ms; p = 8.19e-6). **B**. Same as in A but for single trial delta power versus pupil area. Baseline activity was again higher and response latencies faster on the wheel (wheel: 12.3 ± 4.1 spikes/s; tube: 9.6 ± 4.5 spikes/s; p = 0.0002; wheel: 44 ± 9 ms, tube: 55 ± 10 ms; p = 8.19e-6). **C**. Same as in A, but for single trial NBG power versus. (wheel: 12.3 ± 3.6 spikes/s, tube: 9.5 ± 4.5 spikes/s, p = 4.18e-5; wheel: 45 ± 8 ms, tube: 56 ± 12 ms; p = 1.01e-05). **D**. Same as in A, but for ME versus pupil area. Baseline remained higher and latencies faster on the wheel (wheel: 12.3 ± 4.1 spikes/s; tube: 9.6 ± 4.5 spikes/s;; p = 0.0002; wheel: 44 ± 8 ms, tube: 55 ± 11 ms; p = 1.98e-5).

**Supplementary Figure 5.**
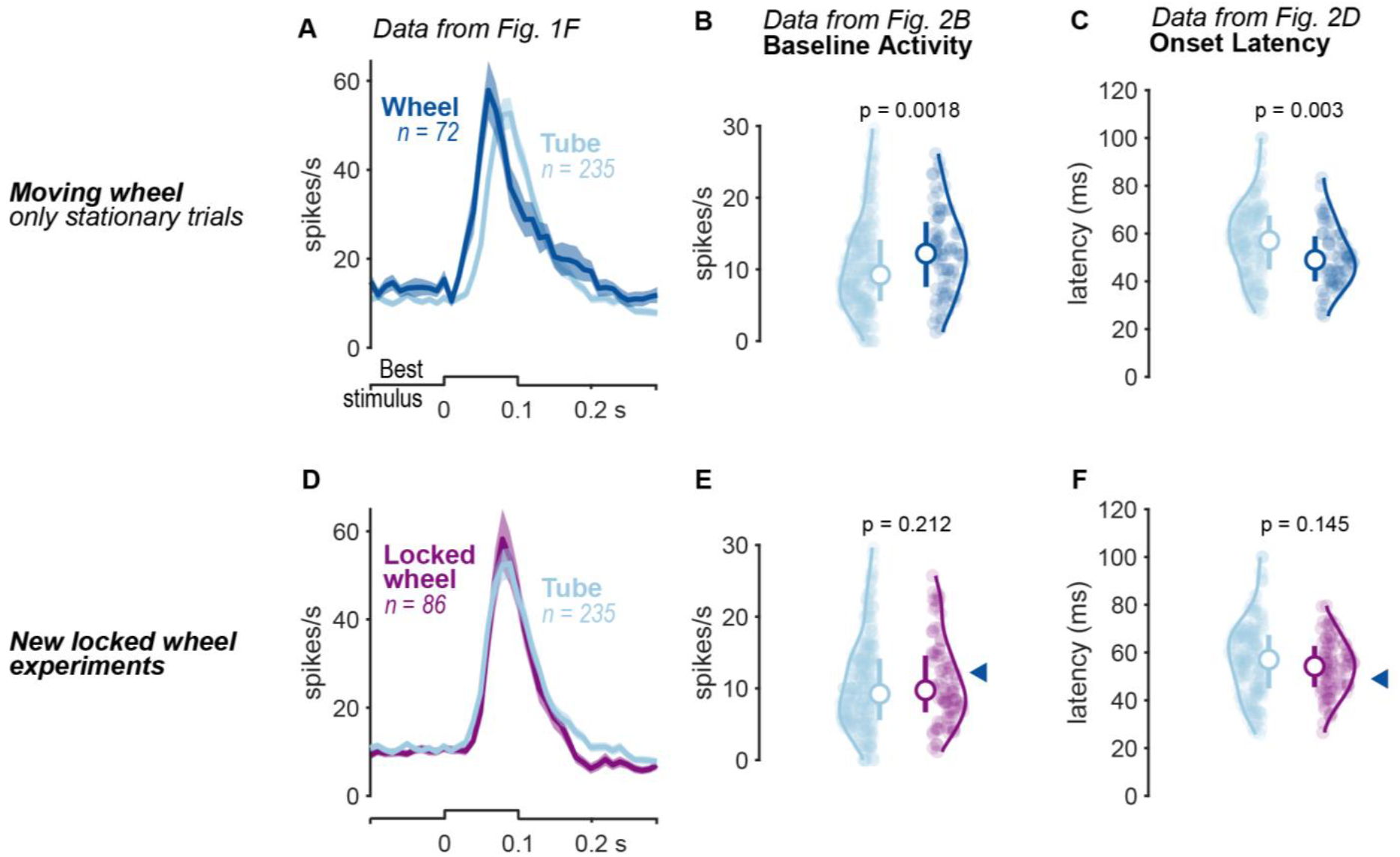
LGN activity on a locked wheel is most similar to the tube. **A**. Data from Figure 1F. dLGN neuron spike responses to the preferred stimulus in the receptive field across contexts (tube, n = 235 neurons; wheel n = 72 neurons). **B**. Data from Figure 2B. Baseline activity was significantly elevated in the wheel context (wheel: 11.4 ± 4.1 spikes/s, n = 107; tube: 8.9 ± 3.4 spikes/s, n=276; *p* = 0.0018; Wilcoxon one-sided rank-sum test, Median ± IQR/2; same statistics throughout figure). Baseline calculated in 0.1s window prior to stimulus onset. **C**. Data from Figure 2D. Onset latency significantly faster on the wheel (wheel: 47 ± 19 ms, tube: 57 ± 24 ms; *p* = 0.003). **D**. New experiments on a locked wheel (purple; n=86 neurons, preventing any possibility of locomotion) show that the LGN population response overlaps responses in the tube (light blue, n=235 neurons). **E**. Baseline activity was not signifcantly different on locked wheel versus tube (tube: 8.9 ± 3.4 spikes/s, n=235; locked wheel: 9.7 ± 7.9 spikes/s, n=86, p = 0.212, Wilcoxon one-sided rank-sum test, Median ± IQR/2; same statistics throughout figure), **F**. Visual onset latency was not signifcantly different on locked wheel versus tube (tube: 57 ± 24 ms; locked wheel: 54 ± 24 ms; p = 0.145).

**Supplementary Figure 6.**
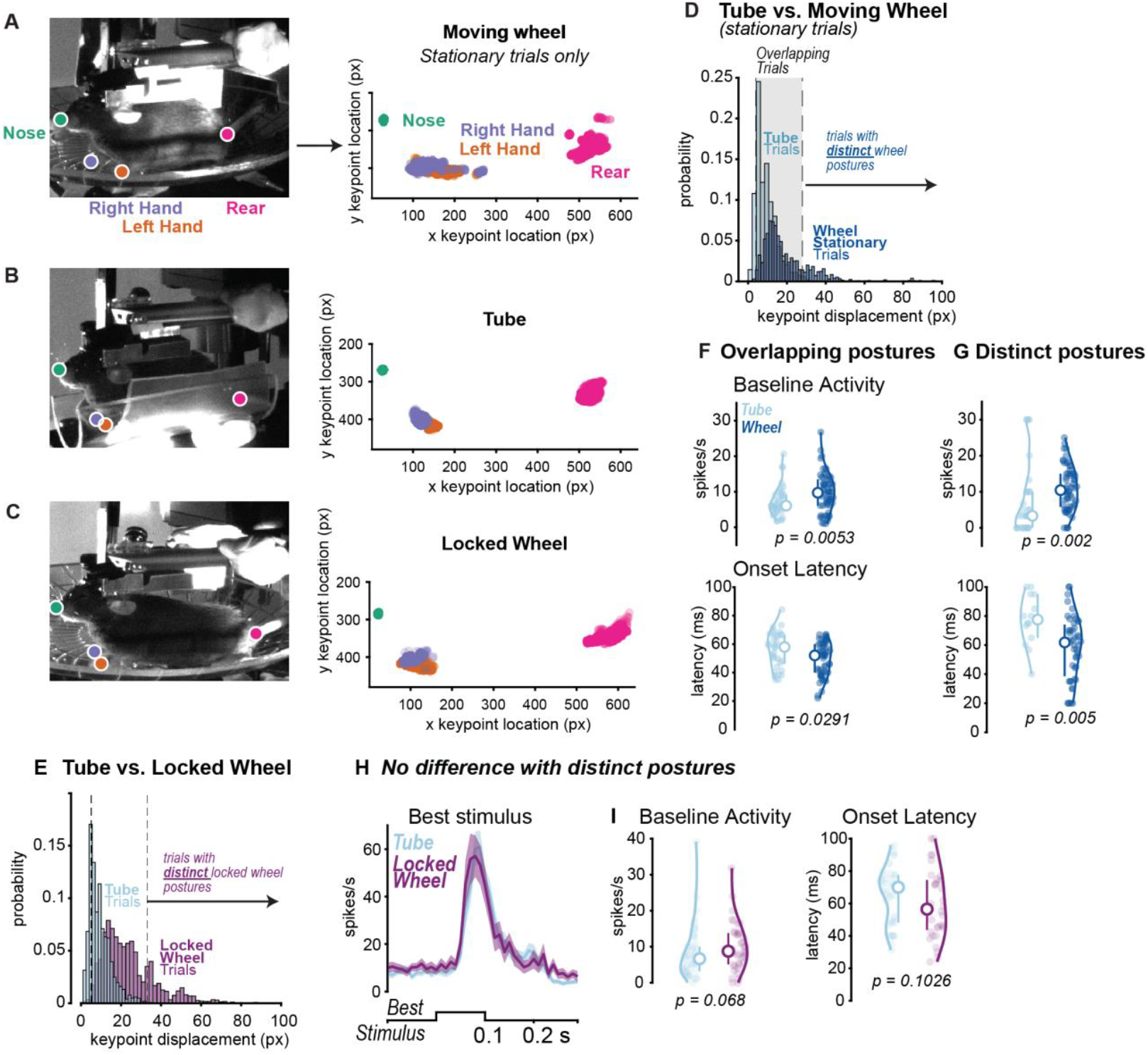
Postural differences do not explain contextual effects on baseline activity and timing. **A**. Moving wheel context. High-speed full body video of mice in either the moving wheel (top; stationary trials) or tube context. Example frames with keypoints labeled at the nose (green), right paw (purple), left paw (orange), and rear (magenta). Keypoint distributions across all frames (22k; 12.3 mins; N = 2 mice. Distribution spread shows (x,y) displacement of that keypoint, due to postural adjustments. Note that nose keypoint (green) remains still and has a nearly identical (x,y) position across both contexts (identical head fixation position). Wheel context was limited to stationary trials (no movement for >2 s). **B**. Same as in A, but for tube context. **C**. Same as in A, but for locked wheel context. **D**. At matched arousal levels (as in Fig. 1E), keypoint displacement was plotted in tube (light blue, 3120 trials) and wheel (dark blue, 3721 trials). Overlapping postures were defined by the median +/-mad of the summed histograms. Non-overlapping postures were set outside of this range. **E**. At matched arousal levels (as in Fig. 1E), keypoint displacement was plotted in tube (light blue, 3120 trials) and locked wheel (purple, 4450 trials). **F**. For trials of overlapping keypoint displacement baseline activity (*left*; wheel: 10.0 ± 3.9 spikes/s, n = 63; tube: 7.1 ± 2.4 spikes/s, n=39; *p* = 0.0053; Wilcoxon one-sided rank-sum test, Median ± IQR/2; same statistics throughout figure) and onset latency (*right*; wheel: 51 ± 10 ms, n = 63; Tube 58 ms ± 10 ms, p = 0.0291) significantly different between the two contexts. 1899/3120 overlapping tube trials, 2276/3721 overlapping wheel trials. **G**. For trials of distinct postures with non-overlapping keypoint displacement, baseline activity (*left*; wheel: 11.0 ± 4.9 spikes/s, n = 47; tube: 3.33 ± 5.4 spikes/s, n=14; *p* = 0.0053) and onset latency (*right*; tube: 77 ± 15 ms, n = 14; wheel 60 ms ± 18 ms, n = 47; p = 0.005) significantly different between the two contexts. 137/3120 non-overlapping tube trials, 621/3721 non-overlapping wheel trials. **H**. dLGN spiking response on trials with distinct postures are similar across locked wheel (n = 53 units) and tube (n=39 units). **I**. On trials with distinct postures with non-overlapping keypoint displacement, baseline activity between tube and locked wheel context were not significantly different (wheel lock: 6.7 ± 4.1 spikes/s, n = 53; tube: 8.8 ± 3.4 spikes/s, n=39; *p* = 0.0681; Wilcoxon one-sided rank-sum test, Median ± IQR/2; same statistics throughout figure). Onset latency between tube and locked wheel context were not significantly different (locked wheel: 55 ± 15 ms; tube 70 ms ± 14 ms, p = 0.1026). 603/3120 distinct tube trials, 976/4550 distinct locked wheel trials.

**Supplementary Figure 7.**
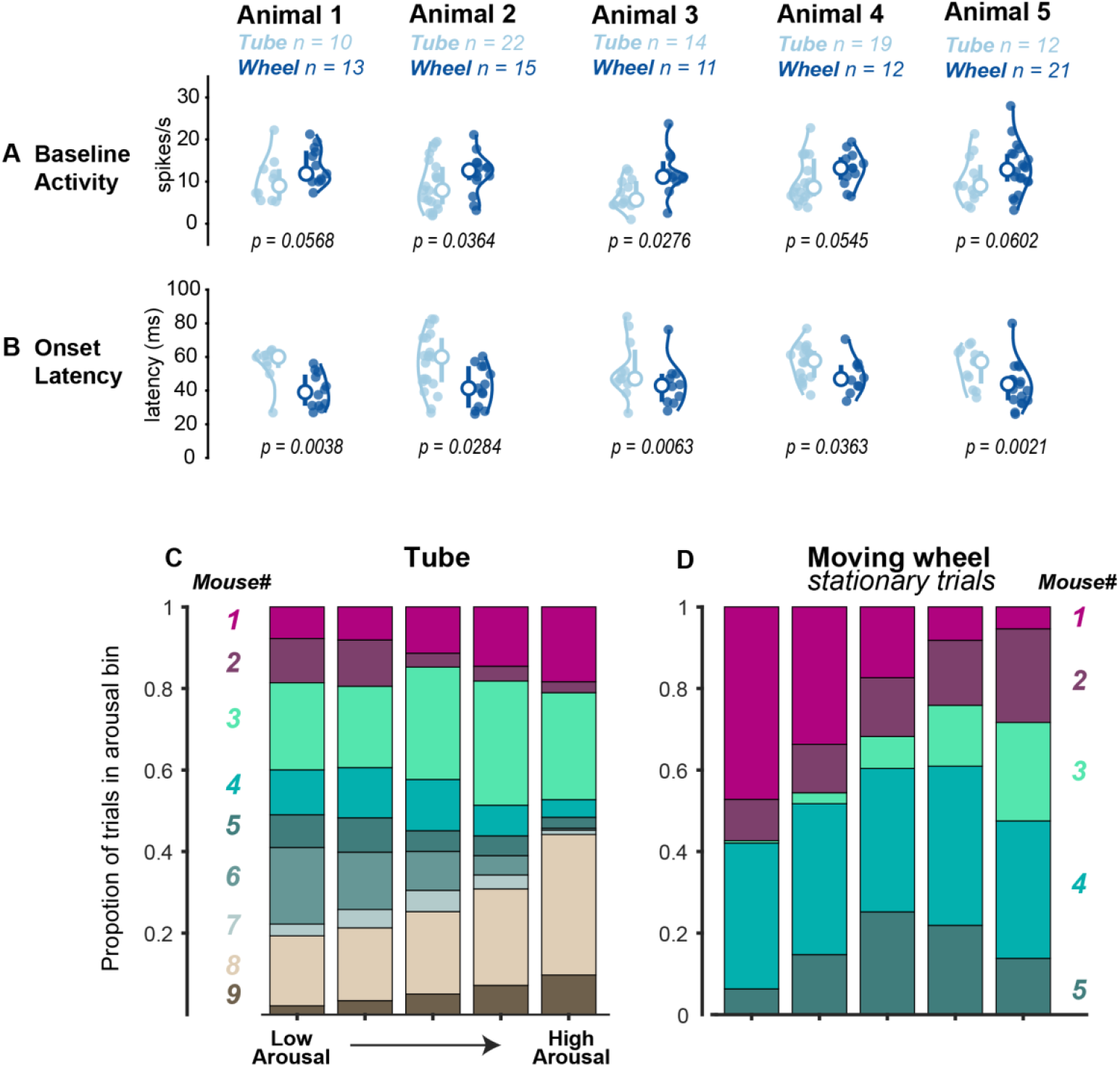
Consistent differences between tube and wheel in baseline activity and onset latency across individual mice. **A**. Baseline activity was significantly elevated in the wheel context relative to the tube in 2/5 mice recorded in both contexts. Statistics for each animal in Supplementary Table 1. **B**. Onset latency was significantly faster in the wheel context relative to the tube in 5/5 mice recorded in both contexts. Statistics for each animal in Supplementary Table 1. **C**. All N = 9 mice recorded from in the tube context contribute data to every arousal bin defined in Figure 3A. **D**. Same as in C, but for the N = 5 mice recorded in the wheel context.

**Supplementary Table 1.** Statistics related to Supplementary Figure 7 for each animal.

| Animal | Tube units | Tube Spont.<br>Med±IQR/2<br>(sp/s) | Tube Onset<br>Med±IQR/2<br>(ms) | Wheel units | Wheel Spont.<br>Med±IQR/<br>2<br>(sp/s) | Wheel Onset<br>Med±IQR/<br>2<br>(ms) | Spont.<br>p-val | Onset<br>p-val |
| --- | --- | --- | --- | --- | --- | --- | --- | --- |
| 1 | 10 | 9.0±3.8 | 60±4 | 13 | 11.9±3.6 | 39±9 | 0.0568 | 0.0038 |
| 2 | 22 | 7.9±4.4 | 60±10 | 15 | 13.3±2.1 | 41±12 | 0.0364 | 0.0072 |
| 3 | 14 | 6.6±2.8 | 48±10 | 11 | 11.2±2.3 | 44±8 | 0.0276 | 0.0963 |
| 4 | 19 | 8.6±4.5 | 58±7 | 12 | 13.2±2.7 | 47±6 | 0.0545 | 0.0230 |
| 5 | 12 | 9.0±3.8 | 57±10 | 21 | 13.3±3.3 | 44±6 | 0.0602 | 0.0063 |

